# Real versus virtual faces: Differentiating neural responses to emotional expressions

**DOI:** 10.64898/2026.04.29.720690

**Authors:** Dylan Rapanan, Steven R. Livingstone, Zedd Whitaker, Ryan Stevenson, Bobby Stojanoski

**Affiliations:** Department of Computer Science, Ontario Tech University, Oshawa, ON, Canada; Department of Psychology, Ontario Tech University, Oshawa, ON, Canada; Department of Psychology, Western University, London Ontario; The Center for Advanced Interdisciplinary Research (CeNIIs) at Ss. Cyril and Methodius University, Skopje North Macedonia

**Keywords:** Virtual faces, Emotion perception, Functional near-infrared spectroscopy (fNIRS), Functional connectivity, Brain networks

## Abstract

As avatars become commonplace, understanding how the brain processes emotion expressed by virtual faces is critical. Across two experiments, we compared behavioral and neural responses to real and virtual faces expressing seven emotions (anger, disgust, fear, joy, sadness, surprise, neutral). In Experiment 1 (n=61), participants rated the similarity of paired real and virtual faces. Expressions conveying the same emotion were rated as highly similar across face types, whereas mismatched emotions yielded substantially lower similarity ratings, indicating perceived emotional meaning was preserved despite differences in face realism. In Experiment 2 (n=91), functional near-infrared spectroscopy was used to measure brain activity while participants viewed the same stimuli. Functional connectivity analyses revealed network level differences between face type and emotion across the brain. Real faces elicited stronger connectivity patterns across frontal, central-temporal, and parietal regions, whereas high-arousal emotions (fear, anger, and joy) were associated with a broader network engagement than other expressions. Complementary general-linear-model analyses revealed increased activation was limited to visual areas for virtual faces, surprise, and neutral expressions. Our results suggest face-type processing occurs in early visual areas, while emotion-type are distinguished by distinct patterns of network level connectivity across the brain for real and virtual faces.

## Introduction

Our faces are powerful social signals. From a single glance, observers can infer our identity, intentions, and our emotional state (Oosterhof & Todorov, 2008; Snoek et al., 2023). Emotional expressions are central to social interactions (Hadley et al., 2022). Most research on emotional face processing has used photographs of real human faces. However, the environment in which faces are encountered is rapidly evolving. Increasingly, social interaction occurs with digitally rendered agents - avatars embedded in video games, virtual reality, telehealth platforms, customer-service interfaces, social robotics, and emerging AI-driven systems (Garety et al., 2024; Kyrlitsias & Michael-Grigoriou, 2022; Lolansen et al., 2025). As avatars and agents become integrated into everyday communication, a fundamental question arises: does the human brain process emotional expressions conveyed by virtual faces in the same manner as those conveyed by biological humans?

Current neuroscientific models (Lindquist et al., 2012; Pessoa, 2017; Satpute & Lindquist, 2019) emphasize that emotional face perception is supported by distributed cortical representations rather than discrete “emotion centers.” Multivariate fMRI studies show that emotional categories can be decoded from activation patterns across occipital, temporal, and frontal cortices (Kragel & LaBar, 2016; Saarimäki et al., 2016). Network-based analyses further indicate that emotion perception modulates inter-regional functional connectivity, reflecting integration of affective and social-cognitive processes (Bayet et al., 2021; Bendall et al., 2016; Westgarth et al., 2021). However, these models are grounded almost exclusively in responses to photographs of real human faces. Whether the same distributed architecture is engaged when emotional information is conveyed by virtual faces is unknown.

A growing body of work has examined our responses to virtual faces. Behavioral studies indicate that anatomically informed avatars can convey emotional expressions with moderate-to-high accuracy (Calbi et al., 2017; García et al., 2020; Hortensius et al., 2018). Neurophysiological findings however are mixed. EEG studies show differences in theta and alpha activity between avatars and real faces (Kegel et al., 2020;Sollfrank et al., 2021). fMRI studies suggest that perceived realism modulates activation in face-sensitive regions and social-evaluative networks (de Borst & de Gelder, 2015; Schindler et al., 2017). However, most prior work typically employed limited emotion sets, small stimulus samples, or single avatar identities, constraining generalizability. Collectively, this leaves an open question: do differences in neural processing of real and virtual faces emerge primarily at early perceptual stages, or at the level of distributed network coordination for higher-order emotional evaluation?

Across two experiments, we explored participants perceptual and neural responses to matched expressions from diverse real and virtual faces. In Experiment 1, we established perceptual parity by matching an open-access virtual dataset to an existing real-face dataset. In Experiment 2, we presented these same faces to participants while recording neural responses using functional near-infrared spectroscopy (fNIRS). By integrating activation and network-level analyses, we aim to determine whether emotional expressions conveyed by virtual faces recruit neural systems comparable to those engaged by real faces, or whether differences emerge that reflect the perceptual and social status of synthetic faces. Specifically, we hypothesised that 1) virtual and real faces would produce distinct activation and connectivity profiles, and 2) different emotional expressions would evoke unique neural signatures.

## Experiment 1

In Experiment 1, we established perceptual equivalence for the real-virtual comparison used in Experiment 2. This required a virtual-face dataset that was demographically diverse, contained multiple emotions per model, and supported matching across face identities. Existing Action Unit-driven systems are designed primarily for photo-realistic faces rather than clearly virtual avatars (Krumhuber et al., 2012), other virtual-face stimuli are not generally available as reusable open-access datasets (Joyal et al., 2014), and generative AI tools cannot reliably produce a fixed identity with a series of different expressions. We selected the UIBVFED (Oliver & Amengual Alcover, 2020), an open-access dataset of 20 demographically varied virtual characters expressing 32 discrete emotions using blendshape-based modelling that approximate Facial Action Coding System (FACS) principles (Lewis et al., 2014). These faces were paired with a validated set of faces (RADIATE; Conley et al., 2018) that were matched for sex and race/ethnicity expressing overlapping emotion categories.

Participants first completed an emotion-recognition task to test whether the selected UIBVFED expressions conveyed their intended emotions above chance. Participants then completed a similarity-rating task in which real and virtual faces were paired under conditions that varied face-type congruence and emotional-category congruence. We predicted the highest similarity ratings for pairs matched on both identity and emotion, intermediate ratings for pairs matched on emotion but differing in face type, and the lowest ratings for pairs matched on identity but differing in emotion.

### Methods

#### Participants

Sixty-one students from Ontario Tech University participated in the study (20 men and 41 women, M = 20.84, SD = 3.25, range = 17 to 38). The experiment took approximately 40 minutes and participants received course credit. The study was approved by the Ontario Tech University Research Ethics Board (REB: 17656).

#### Stimuli

One hundred and forty facial images from the RADIATE and UIBVFED datasets were used (Conley et al., 2018; Oliver & Amengual Alcover, 2020). The RADIATE contains perceptually validated images of racially and ethnically diverse participants, aged 18-30 years old, each expressing 16 different emotions. The UIBVFED dataset contains a set of 20 virtual characters that are also ethnically diverse, aged 20-80 years old, forming 32 distinct facial expressions. Ten adult models (5 men, 5 women) from each dataset were identified and matched between-sets on face shape, sex, skin tone, and hair colour. Two raters independently matched model identities between the UIBVFED and RADIATE sets, with models achieving the highest inter-agreement selected. Images of selected models expressing seven emotions were then selected (Neutral, Joy, Sad, Angry, Fearful, Surprise, Disgust). Selected emotions closely align with basic emotions (Ekman & Cordaro, 2011; Ekman & Friesen, 1971) commonly used in emotion datasets and experiments (Livingstone & Russo, 2018). The full set of images is presented in Supplemental 1.

##### Procedure and design

The experiment employed a two-stage rating task. In stage one, participants identified the emotion portrayed by virtual avatars from the UIBVFED with a forced-choice rating task, shown in Figure 1a. For each trial, a fixation cross was shown for one second, then a photo of an avatar expressing an emotion was displayed in the center of the screen. Participants were then asked to identify the emotion being expressed by selecting one of the seven emotions. Only the virtual avatars from the UIBVFED stimulus set were used in this emotion recognition task, as the RADIATE dataset was previously validated for emotion recognition.

**Figure 1:**
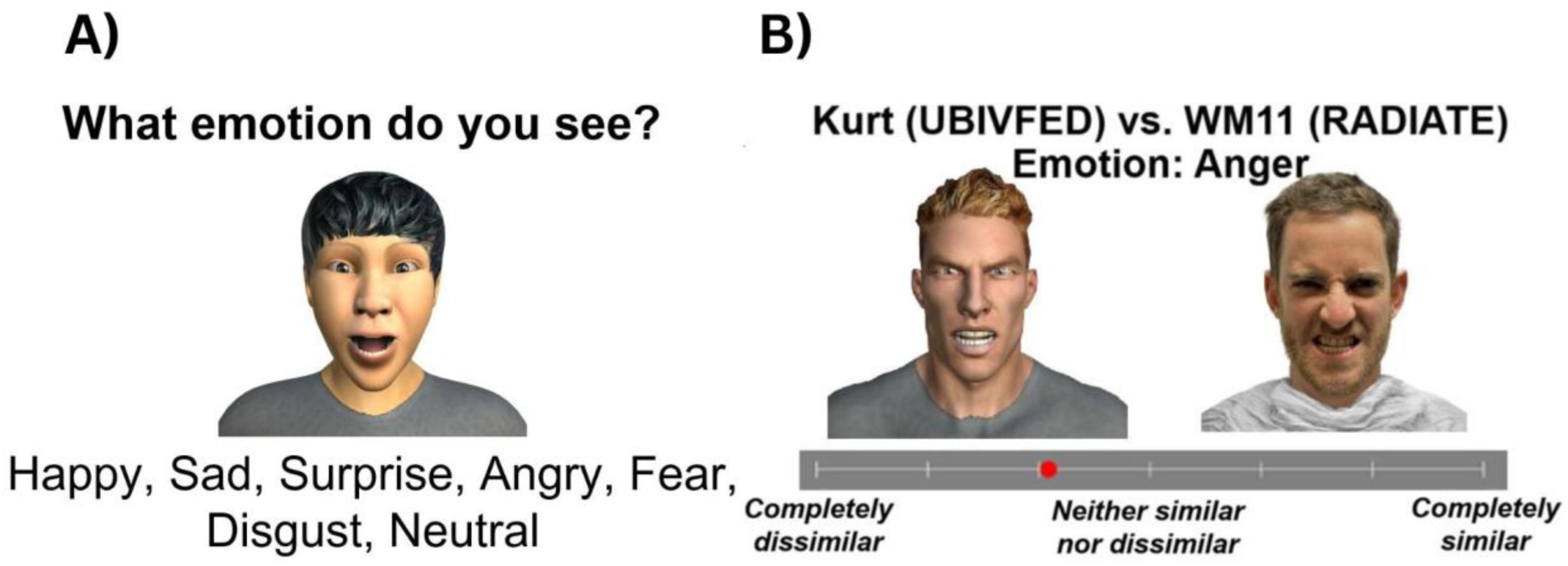
Validation task showing a) Example trial from the emotion recognition task using virtual avatars from the UIBVFED stimulus set. b) Example trial from the similarity matching task, CFIE condition.

In stage two, participants rated expression similarity of side-by-side real and virtual models, as shown in Figure 1b. Participants were instructed to rate the similarity of the expressions displayed on the faces, not of the faces themselves. Participants were first shown a fixation cross for one second. Following this, a pair of randomly selected photos, one virtual and one real, displayed side-by-side on screen with a slider bar underneath to make the similarity rating. There were three conditions in the similarity matching task: 1) Congruent-Face-Congruent-Emotion (CFCE) condition, in which photos were matched on both model-face and expression similarity, 2) Incongruent-Face-Congruent-Emotion (IFCE), mismatched on model-face similarity and matched on expression similarity, and 3) Congruent-Face-Incongruent-Emotion (CFIE), matched on model-face similarity and mismatched on expression similarity.

We used a full-factorial design for our similarity task, consisting of within-subject factors: Congruence (3 levels: CFCE, IFCE, CFIE) × Emotion (7 levels: Neutral, Happy, Sad, Angry, Fearful, Surprise, Disgust) × Model (5) × Sex (2). The 140 trials were presented in random order.

#### Analyses

Statistical analyses were conducted in R (R Core Team, 2026). Participant responses were coded as correct (1) when the rater’s label matched the stimulus’ target label, otherwise incorrect (0). Accuracy was aggregated within each rater and emotion as the proportion correct across trials of that emotion. For each emotion, we tested whether mean per-rater accuracy exceeded chance performance (*p*_0_ = ^1^⁄_7_) using one-sided one-sample t-tests with Holm adjustment. Mean accuracies and 95% confidence intervals are reported for each emotion

Similarity ratings (1-7) were analysed using a Gaussian linear mixed-effect model fit with restricted maximum likelihood in R. Fixed effects were Congruence × Emotion, with random intercepts and by-participant congruence-slopes (1 + congruence | participant). Fixed-effect Type III tests used Kenward–Roger (KR) degrees of freedom; planned contrasts on estimated marginal means (CFCE–IFCE and CFCE–CFIE within each emotion) used KR with Holm adjustment for multiple comparisons. Effect size for the overall model was calculated as conditional coefficients of determination and interpreted using Cohen’s (2013) guidelines. Additional details on model building and analyses are presented in Supplementary 1.

### Results

Emotion identification accuracy was significantly above chance for all emotions (all *p* < 2.23 × 10^−4^). These results confirmed selectMean ratings, along with those for matched RADIATE stimuli, are presented in Table 1. Full test results are reported in Supplemental 1.

**Table 1:** Emotion validation accuracy ratings with chance accuracy one-sided one-sample t-tests (Holm-corrected), and RADIATE ratings.

| Emotion | UBIVFED |  |  |  | RADIATE | Mean |
| --- | --- | --- | --- | --- | --- | --- |
|  | M (SD) | t (df) | p | 95% CI | M (SD) |  |
| Neutral | .89 (.14) | 42.7 (60) | < .0001 | [.86, .93] | .89 (.13) | .89 |
| Happy | .59 (.24) | 14.4 (60) | < .0001 | [.53, .65] | .8 (.2) | .7 |
| Sad | .89 (.12) | 48.0 (60) | < .0001 | [.85, .92] | .7 (.26) | .8 |
| Angry | .5 (.2) | 13.9 (60) | < .0001 | [.45, .55] | .69 (.24) | .6 |
| Fear | .24 (.2) | 3.72 (60) | = 0.0002 | [.19, .29] | .48 (.22) | .36 |
| Surprise | .83 (.17) | 32.1 (60) | < .0001 | [.78, .87] | .84 (.14) | .83 |
| Disgust | .69 (.26) | 16.6 (60) | < .0001 | [.62, .76] | .56 (.23) | .62 |
| Mean | .66 (.08) |  |  |  | .71 (.19) | .68 |

Expression similarity ratings are presented in Figure 2. The model explained a substantial proportion of variance in similarity ratings (*R*^2^=0.49, 95% CI [0.44, 0.53], ICC = 0.18). There was a main effect of Congruence (*F*(2,59.01) = 330.67, *p* < 0.0001). Planned contrasts showed CFCE > IFCE (Δ = 0.32, SE = 0.07, t(2,60) =4.46, p < 0.0001, 95% CI [0.141, 0.491]), CFCE > CFIE (Δ = 2.89, SE = 0.13, t(2,60) = 21.99, p < 0.0001, 95% CI [2.57, 3.22]); and IFCE > CFIE (Δ = 2.58, SE = 0.17, t(2,60) = 15.03, p < 0.0001, 95% CI [2.16, 3.0]). The results indicate that congruent face-emotions were rated as most similar. Importantly, incongruent faces with congruent expressions were also rated as significantly more similar than congruent faces with incongruent emotions, supporting our expectations. There was also a main effect of Emotion (*F*(6,12819.04) = 51.76, *p* < 0.0001), and a significant Congruence × Emotion interaction (*F*(12,12819.04) = 16.44, *p* < 0.0001). Planned contrasts confirmed that all Congruence-Emotion pairs were significantly different except Sad presentations for CFCE and IFCE.

**Figure 2:**
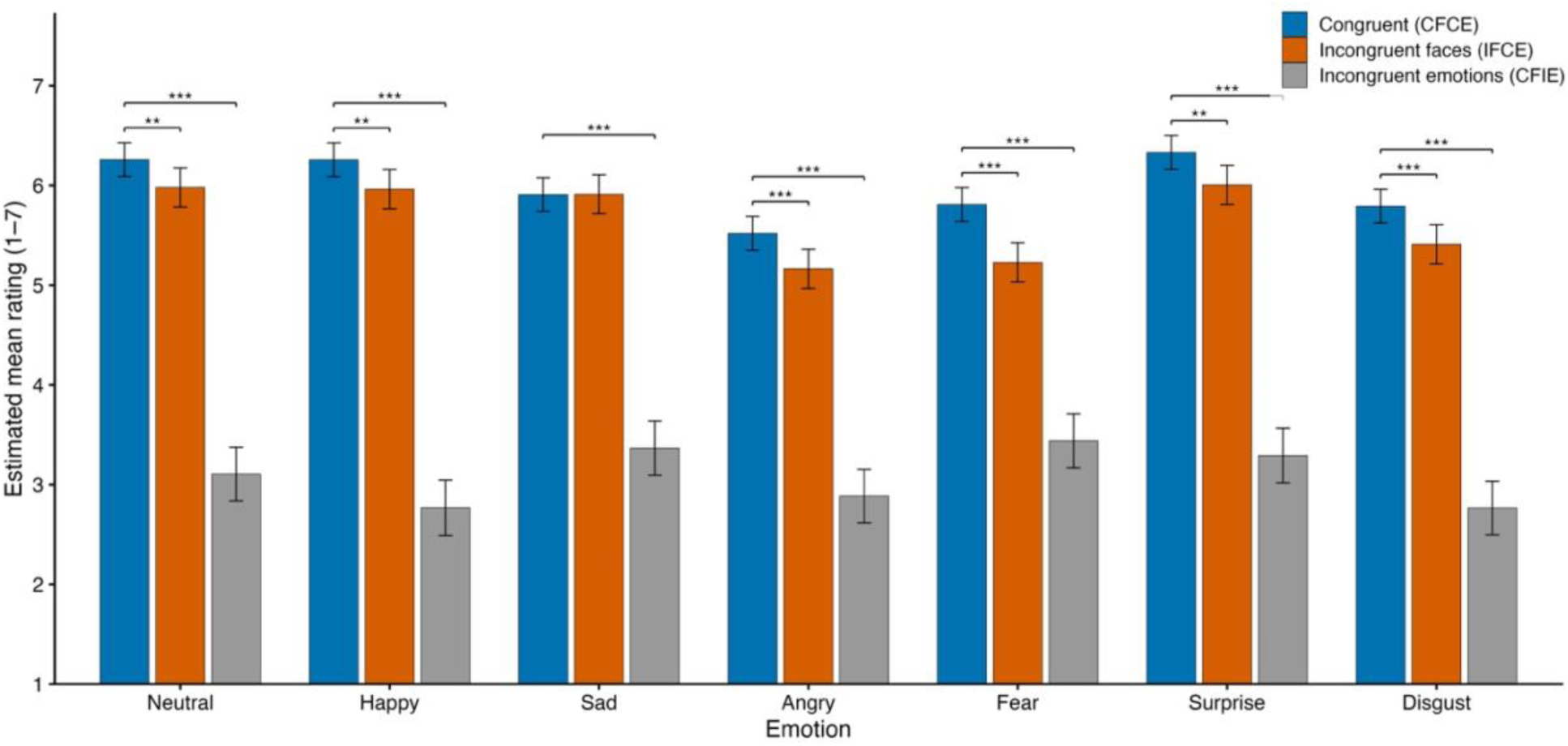
Mean similarity ratings for each emotion across Congruent face and emotion (CFCE), Incongruent faces (IFCE), and Incongruent emotions (CFIE) conditions. Asterisks denote Holm-adjusted one-sided tests against chance: * p <.05, ** p< .01, *** p<.001

### Discussion

Here we evaluated the perceptual validity of selected UIBVFED expressions, and their perceptual correspondence with matched faces from the RADIATE dataset. Participants reliably identified the intended emotion from virtual expressions at rates significantly above chance for all emotion categories. These results indicate that selected UIBVFED expressions reliably conveyed target emotions, supporting their use as experimental stimuli.

Recognition accuracy varied across emotions. Neutral, sad, and surprised expressions were identified with high accuracy, whereas anger and happiness were recognized at more moderate levels, and fear showed the lowest accuracy. Similar variability has been reported in studies of virtual faces, where recognition differences depend on the specific emotion being rendered, with some configurations being more difficult to convey clearly in avatar faces (Dyck et al., 2008; Joyal et al., 2014; Miller et al., 2023).

The similarity-rating task demonstrated that expressions labeled with the same emotion across the real and virtual datasets were perceived as emotionally similar. As predicted, face pairs matched on identity and emotion received the highest similarity ratings (CFCE), followed by pairs matched on emotion but differing in face type (IFCE), with the lowest ratings observed when emotions differed (CFIE). This pattern indicates that similarity judgments were primarily driven by emotion rather than facial identity. Together, these findings establish perceptual congruence between the selected stimuli from the virtual and real datasets and provide an empirical foundation for Experiment 2.

## Experiment 2

### Methods

#### Pre-registration

Experiment 2 was preregistered on the Open Science Framework before data collection began (https://doi.org/10.17605/OSF.IO/76Q8W). Our preregistration includes study design, a prior power analysis, sampling plan, variables, and analysis plan.

#### Participants

A separate sample of 91 participants were recruited. Four participants were removed due to equipment issues. Remaining participants were screened for task attention and fNIRS signal quality. An inclusion criterion of >= 60% accuracy on the behavioral memory task (chance = 50%) excluded one participant. Eighty-seven participants (69 females and 18 males, M = 21.09, SD = 5.91, range = 17 to 51) were analyzed in the behavioral memory task.

Signal quality was computed using Peak Spectral Power (PSP) and Scalp Coupling Index (SCI) (Pollonini et al., 2016). Measures were calculated using a 5-second sliding window across all channels (Bulgarelli et al., 2025; Hernandez & Pollonini, 2020). An inclusion criteria of PSP > 0.1 & SCI > 0.5 excluded thirty-five participants. Fifty-two participants (39 females and 13 males, M = 21.62, SD = 6.67, range = 17 to 51) were included in the final analysis.

Sample size was estimated using a prospective power analysis with the simr package (Green & MacLeod, 2016). A linear mixed-effects model estimated oxygenated HbO with Face-type and Emotion as fixed effects and by-Participant random intercepts. Effects from Gao et al. (2019) for Emotion were used as the basis for simulation estimates. The experiment had more than 80% power [95% CI] to detect effects at 0.05 alpha error probability. Our sample size aligned with comparable studies of emotion processing using fNIRS (Westgarth et al., 2021).

##### Stimuli and apparatus

Stimuli validated in Experiment 1 were reused in Experiment 2. Participants were tested individually in a dedicated, quiet windowless testing room. fNIRS data were collected using two NIRSport2 systems (NIRx Medical Technologies, Berlin, Germany). Each NIRSport2 system was equipped with 16 source and 16 detector optodes, and daisy-chained together for a high density 32×32 optode configuration, as shown in Figure 3. Each neighboring pair of source and detector optode is referred to as a channel, resulting in a total of 103 HbO + 103 HbR channels, with 8 short distance channels (8 HbO, 8 HbR). Mean source-detector separation was 30 mm, and 7mm for short distance channels. Optodes were placed on a flexible fNIRS head cap (NIRScap) 58 cm in circumference. Light was emitted at 760 nm and 850 nm wavelengths, with a mean sampling rate of 6.105 Hz.

**Figure 3:**
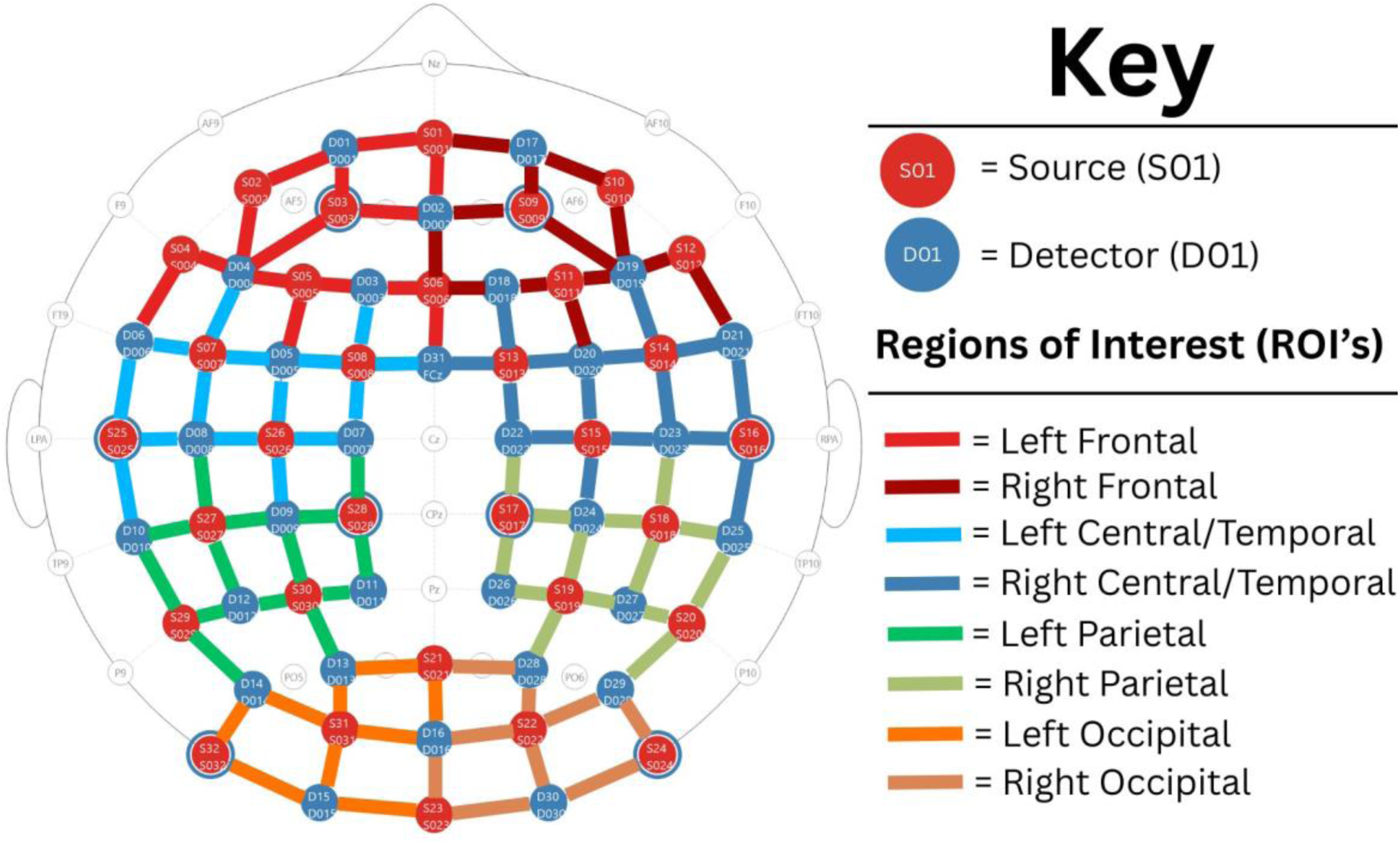
Optode 32×32 montage, coloured to show sources as red vertices, detectors as blue vertices, channels as coloured edges, and short distance detectors as blue-ringed sources (8x). Channels coloured to represent grouped Region of Interest.

#### Design

We used a full-factorial design, consisting of within-subject factors Face-type (2 levels: Real, Virtual) × Emotion (7 levels: Neutral, Joy, Sad, Angry, Fearful, Surprise, Disgust) × Model (4) × Sex (2) × Repetition (4). The 448 images were blocked by Face-type and Emotion, and counterbalanced. Each of the 56 trials (blocks) contained 8 distinct model faces (4 male, 4 female), with leave-one-out for the behavioral memory task.

#### Paradigm

The trial timeline consisted of three main epochs: fixation cross, stimulus block, and memory-task, as shown in Figure 4. Facial images were shown for 1.5 seconds within each block, with a 250-750 ms (M=500 ms) interstimulus interval. In the memory task, participants were presented a probe model image, matched on emotion, and asked if the model was shown in the preceding block. Probe face was present in 50% of trials. Participants were given a break every seven blocks and prompted when ready to continue the experiment. After the experiment was completed, the experimenters re-entered the room, removed the fNIRS cap, and the participant was debriefed. Participation took approximately 35 minutes.

**Figure 4:**
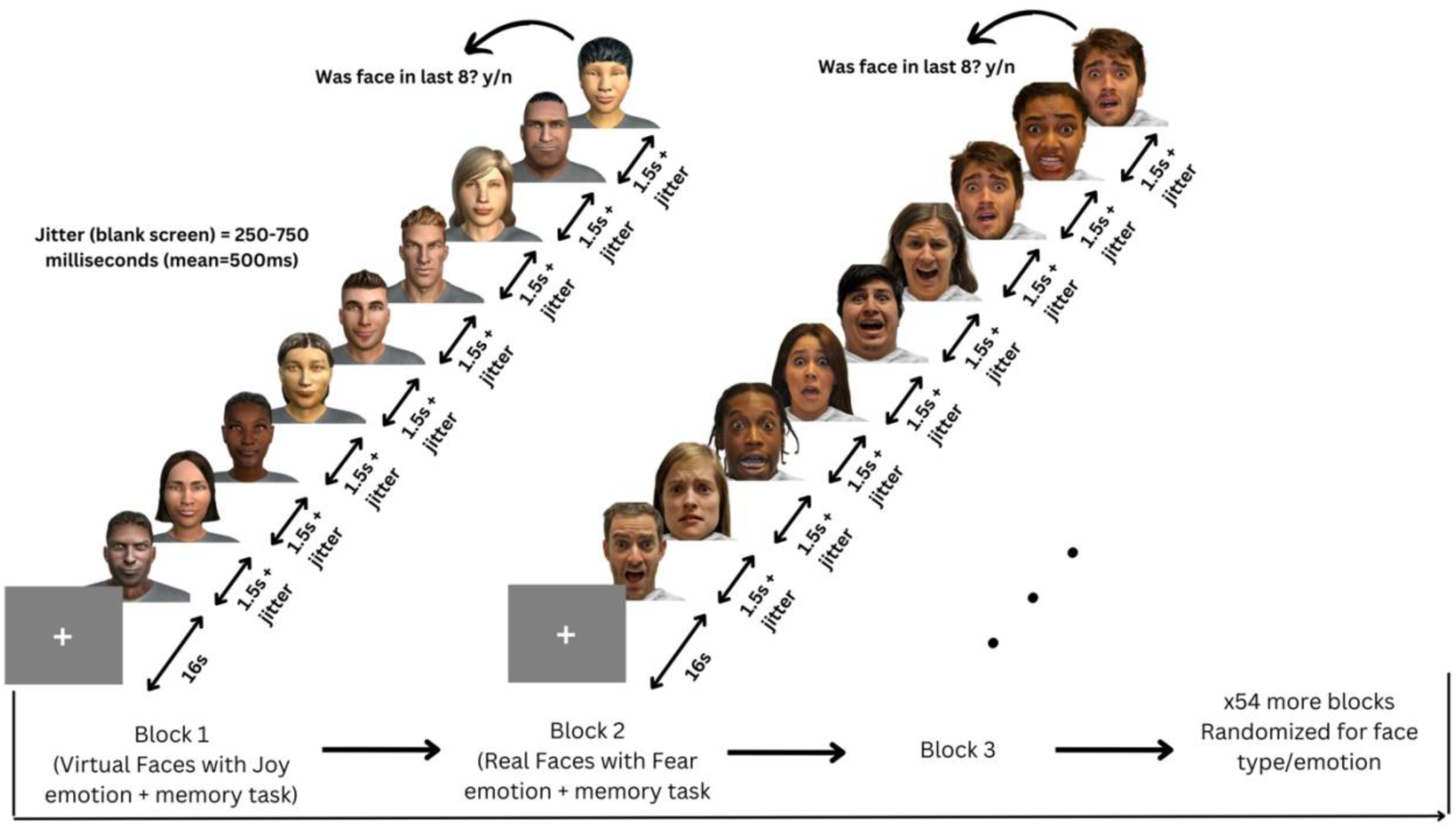
Trial timeline of stimulus presentation. Participants were presented 56 blocks, each containing of 8 distinct facial images. Each Block presented a single face-type and emotion (e.g., Real-Joy faces).

#### Analyses

All fNIRS data were preprocessed and analyzed with Python 3.11.9 using MNE (version 1.9.0) (Gramfort et al., 2013) and MNE-NIRS (version 0.7.1) (Luke et al., 2021), which used the Nilearn package (version 0.9.2). Data were analyzed with a General Linear Model (GLM), followed by a Functional Connectivity (FC) analysis. The memory task was analyzed in Python using the statsmodels package (version 0.14.4) (Seabold & Perktold, 2010).

The GLM estimated the observed haemodynamic signal at each channel or Region of Interest (ROI) as a linear combination of task-related regressors convolved with a Hemodynamic Response Function, plus nuisance regressors (e.g., drift) and residual noise. A two-way repeated measures GLM was conducted on participant’s HbT responses, with Face-type (2) and Emotion (7) entered as fixed effects. Pairwise contrasts were computed between conditions of interest. Significance levels were corrected for multiple comparisons with FDR using the Benjamini-Hochberg procedure (Singh & Dan, 2006) with a family-wise error rate of α=0.05.

This analysis characterized the temporal coordination between channels by computing FC matrices using a continuous wavelet transform (CWT). The morlet wavelet was picked as it is well suited for capturing both slow neural rhythms and faster systemic fluctuations in fNIRS data (Reddy et al., 2021). The frequency range was narrowed to five evenly spaced frequences between 0.2-0.5 Hz due to short epoch length, which targets systemic and neurogenic oscillations while avoiding high-frequency noise (Xu et al., 2017). Averaging across the closely spaced frequencies reduced data dimensionality.

To assess differences in FC, paired t-tests were conducted on unique channel pairs. For each mode (Face type/Emotion), and pair of conditions (e.g., Joy vs. Fear), individual-level connectivity matrices were extracted by averaging across epochs to obtain symmetric channel-by-channel coherence matrices. Fisher’s r-to-z transform (atanh) was applied to each matrix element to normalize the data prior to parametric testing. Paired samples t-tests for each unique channel pair (i > j) were then conducted across participants, with p-values FDR-corrected with a family-wise error rate of q=0.05 (Singh & Dan, 2006).

### Results

#### Responses to real and virtual faces

A main effect of Face Type was reported by the GLM, with pairwise contrasts revealing greater activation for virtual faces compared to real faces, shown in Figure 5a. The contrast of real-virtual faces revealed significant differences in functional connectivity across the brain, shown in Figure 5b. Real faces were associated with increased connectivity predominantly between the left-right parietal, left frontal-left parietal, left central/temporal-left parietal, left central/temporal-right parietal, and the right central/temporal to left parietal regions. In contrast, processing of virtual faces was associated with only slightly increased connectivity between the left central/temporal to right frontal region.

**Figure 5:**
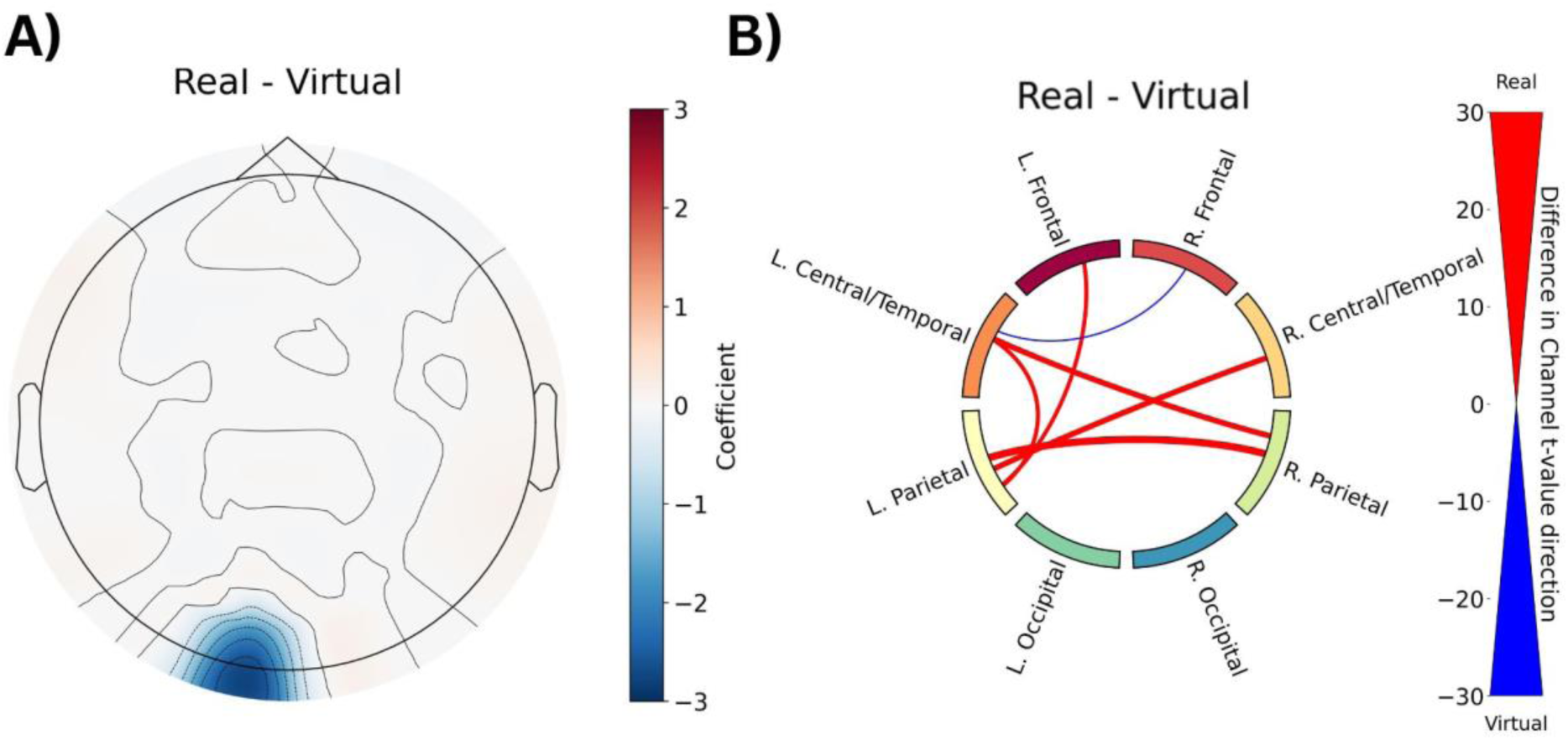
(a) GLM contrast between real and virtual faces. Blue shading indicates stronger activation to virtual faces over real faces (coefficient of the contrast). (b) FC contrast for real and virtual faces. Line thickness represents count of significantly different channel pairs. Line colouring represents t-value sign when real faces had more significant channels between ROI’s (red=positive, blue=negative). Top 15th percentile of connections displayed.

#### Connectivity profiles of facial emotion

To further explore higher level patterns in Emotion, we calculated the ratio of significant channels. Ratios for each emotion-pair were plotted, shown in Figure 6. The ratios were calculated by taking the difference of the count of significant channels where the t-value was positive (red) and negative (blue) and dividing it by the total number of significant channels for that emotion pair. A cluster of emotions with Fear > Anger > Joy connectivity, and Joy, suggests a neural prioritization of faces signalling potential threats, consistent with rapid threat-detection mechanisms. Other emotions (Neutral, Sadness, Surprise, Disgust) had lower connectivity compared to Anger, Fear, and Joy, with Disgust having higher connectivity than Neutral.

**Figure 6:**
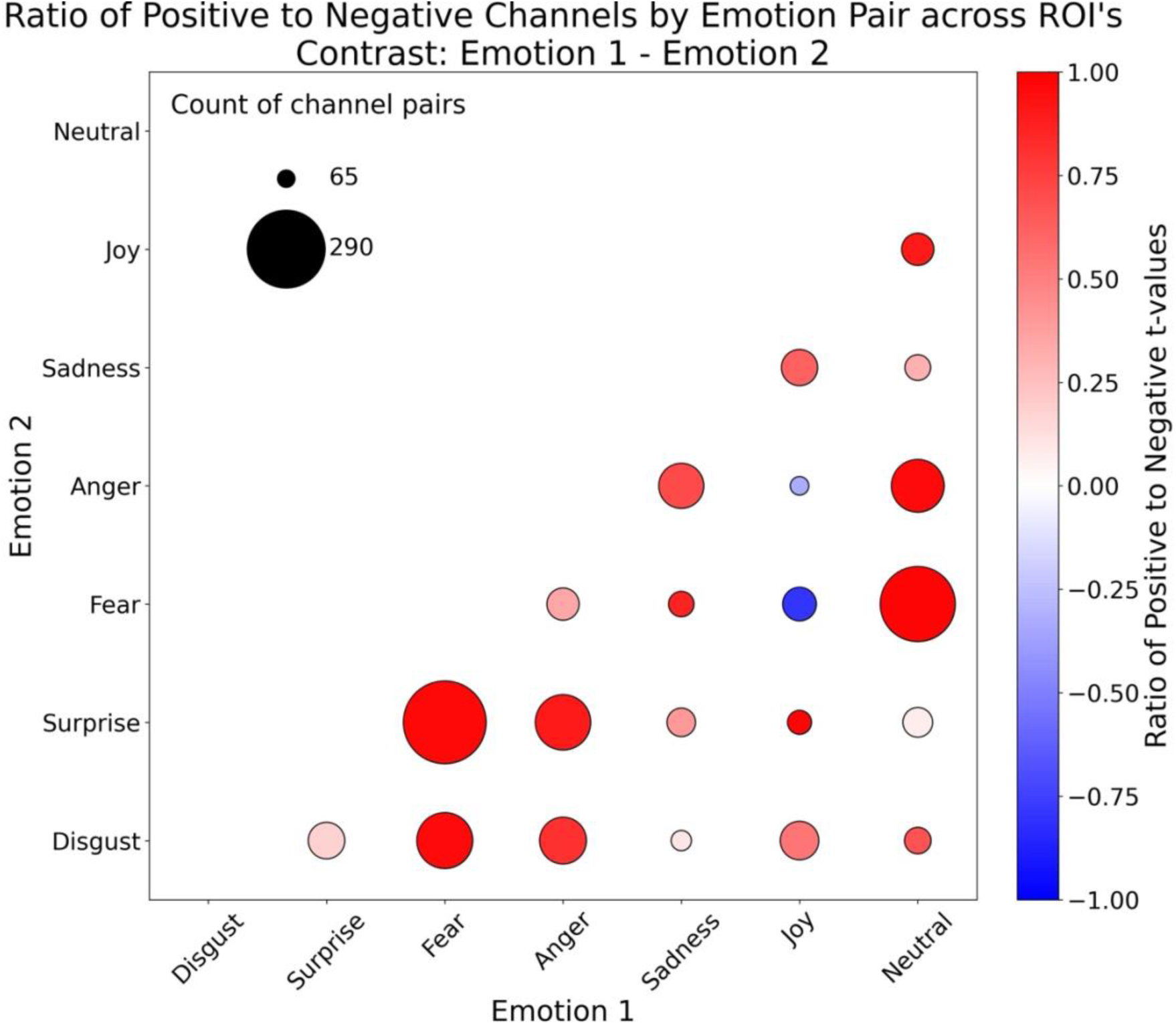
Functional connectivity contrasts of emotion-pairs across ROI’s. Circle diameter represents number of significant channels in emotion-pair tests. Circle colouring represents ratio of significant channel t-value signs (red=positive, blue=negative).

Next, we explored regional patterns of FC. Counts of significant channel pairs per ROI revealed that the left central/temporal, parietal, and right central/temporal regions produced the greatest number of significant connections, shown in Figure 7. Results suggest that these regions are more variable in their connectivity patterns across emotions. In contrast, left and right occipital regions produced the fewest number of significant connections, suggesting that the connectivity within and between these regions are relatively similar in response across emotions.

**Figure 7:**
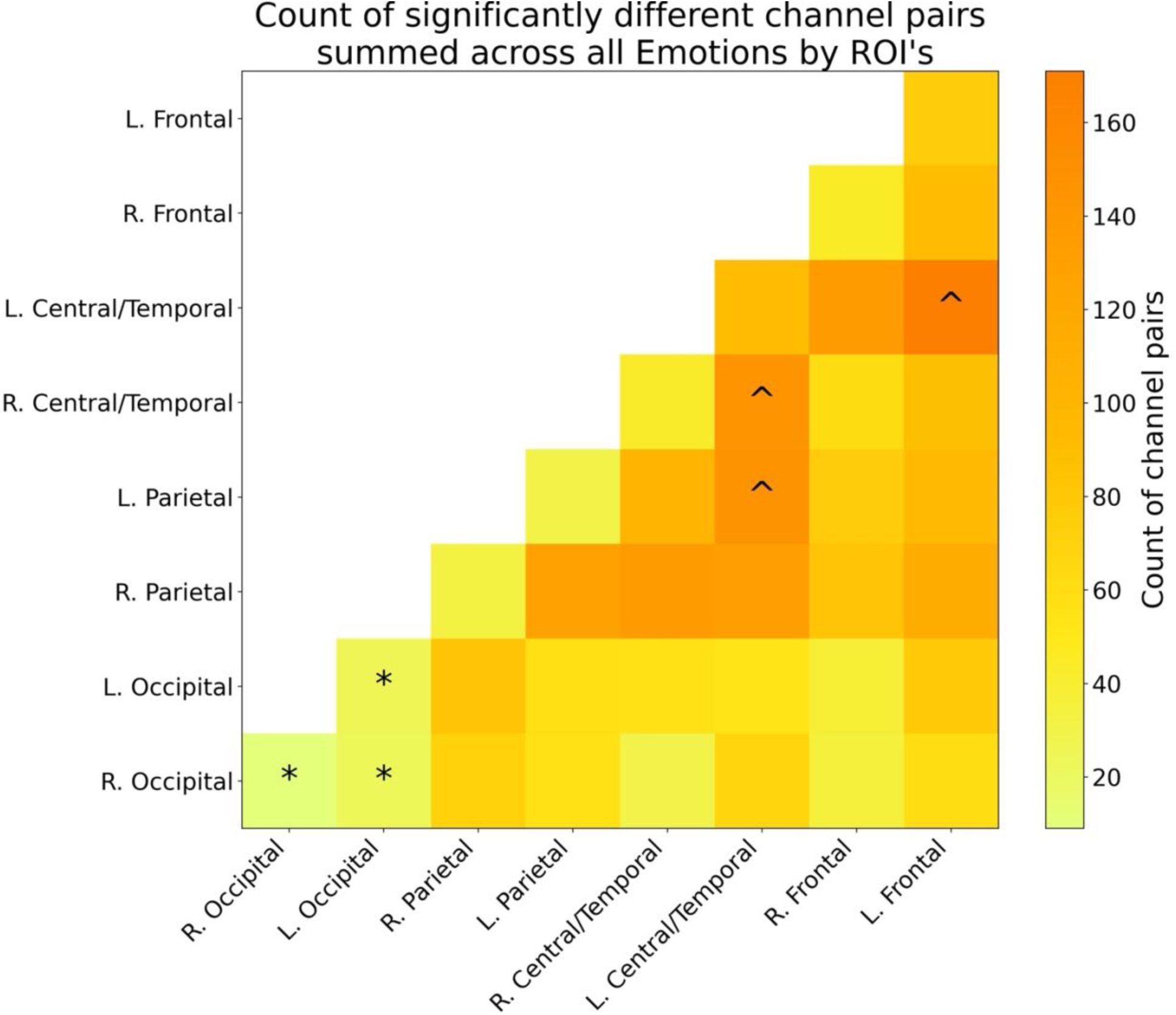
Functional connectivity analysis of number of significantly different channel pairs for each ROI, summed across Emotion. Tile luminance represents number of significant channel pairs (darker=greater, lighter=fewer). ROIs with the three greatest and fewest number of significant channel pairs labelled with ^ and *, respectively.

#### Interaction of face type and emotion

We explored the interaction of Face Type and Emotion by calculating the number of significant channels for each emotion, separately for real and virtual faces, as shown in Figure 8. Joy, Sadness, and Surprise had a greater number of significant channels for real faces compared to virtual faces, while Fear, Anger, Disgust, and Neutral all had a greater number of significant channels for virtual faces compared to real faces. To find which brain regions were driving these differences, we calculated the number of significant channels by Face Type and region, across all Emotions. The occipital areas were found to have a similar number of significant channels for both face types, while the central/temporal and parietal regions had a greater number of significant channels for real faces compared to virtual faces. The frontal regions had a greater number of significant channels for virtual faces compared to real faces.

**Figure 8:**
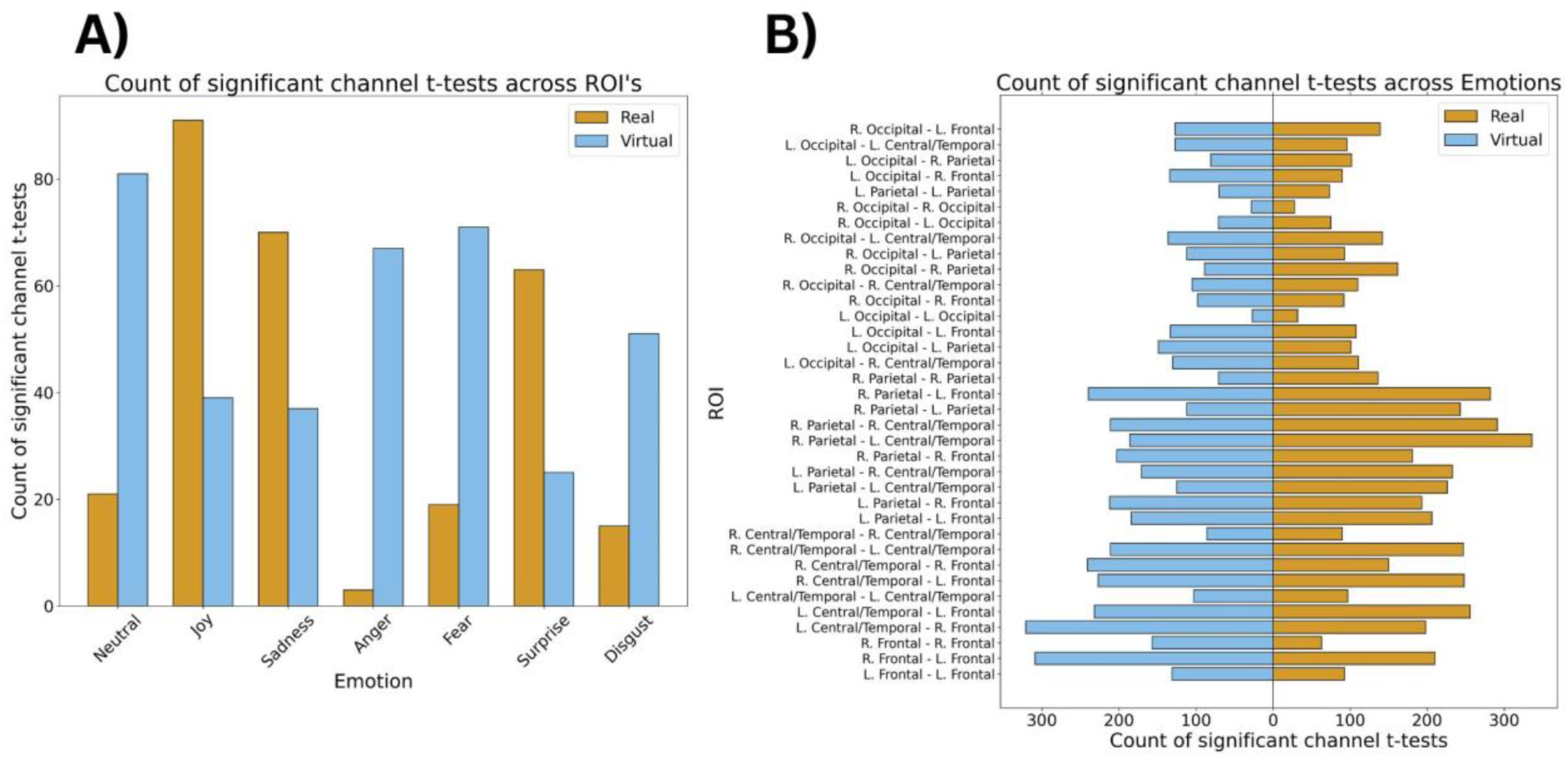
Bar plot summaries of functional connectivity separated by Face Type. Bar height represents the count of significant t-tests, t > 1 for Real and t < −1 for Virtual faces. a) Count of significantly different channel pairs for each emotion. b) Count of significantly different channel pairs for each region, summarized across all emotions.

### Discussion

Distinct neural mechanisms were associated with processing of real and virtual faces. Virtual faces elicited greater activation in left occipital regions compared to real faces, suggesting that face realism modulates early visual processing and likely reflects sensitivity to basic perceptual differences rather than higher-level facial configuration.

In contrast, FC analyses revealed more widespread differences across conditions. Connectivity patterns differentiated both face type and emotion across distributed regions, indicating that network-level dynamics were more sensitive to these factors than localized hemodynamic responses. Interestingly, FC analyses revealed similarity-based clustering whereby fear and anger produced similar patterns of connectivity, but those patterns were quite different than surprise, disgust, and neutral. Conversely, processing surprise, disgust, sadness, joy and neutral expressions produced the most similar patterns of connectivity profiles.

We also observed an interaction between face type and emotion. Real faces showed stronger connectivity for joy, sadness, and surprise, whereas virtual faces showed stronger connectivity for fear, anger, disgust, and neutral expressions. These differences varied by region, with the most prominent differences in frontal, central/temporal and parietal regions, and the fewest difference in visual cortex. These regions are associated with higher-order social cognition and the default mode network, supporting constructionist accounts of emotion that emphasize distributed network interactions favoring real faces and frontal regions favoring virtual faces.

### General Discussion

Across two experiments, we provided converging evidence that emotional expressions on real and virtual faces are perceived as similar yet engage distinct neural networks. While matched real and virtual expressions were rated as emotionally similar, systematic differences in neural responses to those same faces indicated that perceptual similarity does not imply neural equivalence.

At the neural level, differences between face types emerged along two dimensions. Virtual faces elicited greater activation in left occipital region, suggesting face realism modulates visual processing at the earliest stages, whereas real faces were associated with stronger and more widespread functional connectivity across frontal, parietal, and temporal regions. These findings indicate that while both face types are processed as emotional stimuli, they differ in how information is integrated across higher-order cortical networks.

Specifically, virtual faces were associated with greater activity over posterior visual areas. These differences likely reflect early visual areas responding to rudimentary physical differences between virtual and real faces rather than configural face information associated with activity in more anterior regions like the fusiform face area. Despite the study was well-powered, we did not find differences in the fusiform face area, or other more anterior regions. This lack of effects may suggest that despite differences in the basic visual features of between real and virtual faces, neural processing between these types of facial images are mostly the same.

By comparison, emotion-related effects were primarily expressed through functional connectivity rather than activation magnitude. High-arousal emotions (fear, anger, joy) showed stronger and more widespread connectivity than lower-arousal emotions, suggesting that emotional differentiation is organized at the level of distributed neural networks, supporting our first hypothesis. These connectivity differences were concentrated in the central/temporal, parietal, and frontal regions, with relatively minimal differences observed in occipital areas, indicating that higher-order systems may drive variability across emotional conditions.

Experiment 1 provided meaningful context for interpreting these neural findings. By demonstrating that matched real and virtual expressions were perceived as emotionally similar, despite some variability in recognition accuracy, we established that differences observed in Experiment 2 are unlikely to reflect mismatches in perceived emotional meaning. Instead, they likely reflect differences in how the brain processes real and virtual faces. Importantly, similarity ratings were highest when the same emotion was displayed on real and virtual faces matched for perceptual features, while different faces displaying the same emotion, produced comparable ratings, indicating that expressions were well matched on emotional intensity.

In Experiment 2, stronger connections for real faces in frontal, parietal and temporal regions indicated greater integration of higher-order cortical processing areas. Although level of activation in these regions was similar across face types, increased functional coupling reflected distinct neural mechanisms differentiating real and virtual face processing.

Few differences in activation magnitude patterns were observed across emotions. Interestingly, activity in response to surprise and neutral differed from the other emotions, and was restricted to bilateral visual areas. This suggests that most emotions across real and virtual faces produce a similar pattern of activity across the cortex, and surprise and neutral differ based on more rudimentary visual features. In contrast, shared and distinct functional connectivity profiles were observed across the brain for the different emotions. For instance, Anger and Joy also produced notably strong connectivity, clustering with Fear (see Figure 6), suggesting these emotions share common engagement patterns despite their different valence profiles. This clustering aligns with dimensional models of emotion, where high-arousal emotions like Fear, Anger, and Joy engage more extensive and coherent neural networks than their low-arousal counterparts (Ke et al., 2025). The emergence of these two distinct clusters, one characterized by high connectivity (Fear, Anger, Joy) and another by relatively lower connectivity (Disgust, Sadness, Surprise, Neutral), suggests that the brain’s functional architecture may organize emotional face processing along an arousal dimension. This finding is particularly important for understanding how virtual characters and avatars might be designed neural engagement, as high-arousal emotional expressions appear to recruit broader brain networks regardless of face realism.

Interestingly, cortical connectivity patterns reflecting the different clusters of emotions were primarily found in the central/temporal, parietal and frontal regions, with the fewest connectivity differences over occipital areas. In fact, we found only 20-40 significantly different channels in the occipital regions, compared to 100-120 in the central/temporal, parietal and frontal regions. This indicates that unlike face perception, which is driven by early, feature-based aspects of the faces subserved by visual areas such as the occipital and the fusiform face areas, differentiating emotions are associated with distributed network-driven cortical mechanisms that include cortical regions core to the default mode network (DMN). The DMN has been increasingly implicated in social cognition, mentalizing, autobiographical memory, and conceptual integration processes that are central to emotion understanding. Rather than passively “resting,” the DMN supports internally generated models of others’ mental and affective states, integrating perceptual input with prior knowledge, contextual information, and self-referential representations. These results therefore align with constructionist accounts of emotion (Barrett, 2006a; Lindquist & Barrett, 2012; Satpute & Lindquist, 2019; Pessoa, 2017); that emotional expressions are actively constructed through the integration of sensory input with conceptual and prior knowledge.

Functional connectivity between parietal, temporal and frontal regions may instantiate top-down models and highlights that emotional categories emerge from distributed network interactions rather than isolated, domain-specific regions (Barrett, 2006b; Lindquist et al., 2012; Satpute and Lindquist 2019; Pessoa 2017).

Interesting patterns of face-type and emotion were also observed. Joy, Sadness, and Surprise showed greater functional connectivity for real faces, while Fear, Anger, Disgust, and Neutral expressions elicited stronger connectivity for virtual faces. Interestingly, the cortical regions associated with differences in connectivity strength differed for emotions on real and virtual faces. That is, central/temporal and parietal regions exhibited greater connectivity for real faces, whereas frontal regions produced greater connectivity for virtual faces. That we found stronger connectivity in frontal regions for Fear, Anger, and Disgust is consistent with previous work showcasing stronger activation of the prefrontal cortex in response to negative, and high-arousal emotions (Etkin et al., 2011; Tupak et al., 2014), suggesting that in frontal cortex responses to emotions incorporates face realism perhaps because features associated with negative valence are exaggerated or made more salient on virtual faces.

Overall, the results show that while real and virtual faces can convey similar emotional meanings at a perceptual level, they engage partially distinct neural mechanisms. This distinction has implications for understanding how humans process and interact with virtual agents in increasingly digital social environments.

#### Limitations

This study was one of the first to use fNIRS to examine neural responses to real versus virtual emotional faces. However, virtual face images were drawn from a single dataset (Oliver & Amengual Alcover, 2020). While UIBVFED faces were generated based on Facial Action Coding System (Ekman & Friesen, 1978), it is unclear whether emotional expression generated on other virtual faces produce similar patterns of brain activity. Future work should explore a wider range of virtual face styles and realism levels to better understand how these factors influence neural processing and generalize across different virtual face types (Schindler et al., 2017; Thomas et al., 2015).

#### Conclusion

The present study examined how the human brain processes emotional expressions conveyed by real and virtual faces, using functional near-infrared spectroscopy (fNIRS) to assess both regional activation and functional connectivity. Across two experiments, we showed that although real and virtual expressions are perceived as emotionally similar, they are not neurally equivalent. Virtual faces elicited greater activation in the left occipital region, consistent with sensitivity to low-level visual features, whereas real faces were associated with stronger and more widespread connectivity across frontal, parietal, and temporal networks. Emotional differences were expressed primarily in connectivity patterns, with high-arousal emotions (fear, anger, joy) engaging broader network interactions than lower-arousal expressions.

These findings indicate that face realism modulates neural processing at both early perceptual and network levels. While virtual faces are processed as emotional stimuli, they engage partially distinct neural mechanisms, particularly in how information is integrated across distributed cortical regions. The results further suggest that emotional expressions are represented through network-level interactions rather than isolated regional responses.

Together, these findings provide new insight into the neural basis of emotional face perception in increasingly digital environments. Future work should examine how variation in avatar realism, dynamics, and context influences both perceptual and neural responses to emotional expressions.

## Data and code availability

All scripts, data files, and the pre-registration plan are publicly available on OSF https://osf.io/d7bzp/overview?view_only=ff580ecf93204dd7a795881972decb5b.

## AI Usage Disclosure

### Manuscrip

The authors used GPT-5 to edit a final draft of the manuscript for flow, tone, and grammatical correctness. The authors reviewed and edited the content as needed and take full responsibility for the content of the publication.

### Source code

The authors used GPT-5 to edit statistical analysis source code for correctness. The authors reviewed and edited the code as needed, and take full responsibility for the content.

## Acknowledgements

We would also like to thank Dr. Miquel Mascaro-Oliver for sharing the virtual stimuli they created.

## Author contributions

DR involved in design, data collection, and analysis of Exp2, and preparation and review of the manuscript. ZW involved in the design, data collection and analysis of Exp1. RS was involved in the conceptualisation, design and data collection of Exp 1 and discussions about Exp 2. BS was involved in conceptualisation of the project, design of Exp1 and Exp2, analysis of Exp1 and Exp 2, and preparation and review of the manuscript. SRL. was involved in conceptualisation of the project, design of Exp2, analysis of Exp1, and preparation and review of the manuscript. BS and SL contributed equally

## References

Barrett, L. F. (2006a). Are emotions natural kinds? Perspectives on Psychological Science, 1(1), 28–58. 10.1111/j.1745-6916.2006.00003.x

Barrett, L. F. (2006b). Solving the emotion paradox: Categorization and the experience of emotion. Personality and Social Psychology Review, 10(1), 20–46. 10.1207/s15327957pspr1001_2

Bayet, L., Perdue, K. L., Behrendt, H. F., Richards, J. E., Westerlund, A., Cataldo, J. K., & Nelson, C. A., III. (2021). Neural responses to happy, fearful and angry faces of varying identities in 5- and 7-month-old infants. Developmental Cognitive Neuroscience, 47, Article 100882. 10.1016/j.dcn.2020.100882

Bendall, R. C. A., Eachus, P., & Thompson, C. (2016). A brief review of research using near-infrared spectroscopy to measure activation of the prefrontal cortex during emotional processing: The importance of experimental design. Frontiers in Human Neuroscience, 10, Article 529. 10.3389/fnhum.2016.00529

Bulgarelli, C., Blasi, A., McCann, S., Milosavljevic, B., Ghillia, G., Mbye, E., Touray, E., Fadera, T., Acolatse, L., Moore, S. E., Lloyd-Fox, S., Elwell, C. E., Eggebrecht, A. T., & BRIGHT Study Team. (2025). Growth in early infancy drives optimal brain functional connectivity which predicts cognitive flexibility in later childhood [Preprint]. bioRxiv. 10.1101/2024.01.02.573930

Calbi, M., Heimann, K., Barratt, D., Siri, F., Umiltà, M. A., & Gallese, V. (2017). How context influences our perception of emotional faces: A behavioral study on the Kuleshov effect. Frontiers in Psychology, 8, Article 1684. 10.3389/fpsyg.2017.01684

Cohen, J. (2013). Statistical power analysis for the behavioral sciences (2nd ed.). Routledge. 10.4324/9780203771587

Conley, M. I., Dellarco, D. V., Rubien-Thomas, E., Cohen, A. O., Cervera, A., Tottenham, N., & Casey, B. (2018). The racially diverse affective expression (RADIATE) face stimulus set. Psychiatry Research, 270, 1059–1067. 10.1016/j.psychres.2018.04.066

de Borst, A. W., & de Gelder, B. (2015). Is it the real deal? Perception of virtual characters versus humans: An affective cognitive neuroscience perspective. Frontiers in Psychology, 6, Article 576. 10.3389/fpsyg.2015.00576

Dyck, M., Winbeck, M., Leiberg, S., Chen, Y., Gur, R. C., & Mathiak, K. (2008). Recognition profile of emotions in natural and virtual faces. PLOS ONE, 3(11), Article e3628. 10.1371/journal.pone.0003628

Ekman, P., & Cordaro, D. (2011). What is meant by calling emotions basic. Emotion Review, 3(4), 364–370. 10.1177/1754073911410740

Ekman, P., & Friesen, W. V. (1971). Constants across cultures in the face and emotion. Journal of Personality and Social Psychology, 17(2), 124–129. 10.1037/h0030377

Ekman, P., & Friesen, W. V. (1978). Facial action coding system. Consulting Psychologists Press.

Etkin, A., Egner, T., & Kalisch, R. (2011). Emotional processing in anterior cingulate and medial prefrontal cortex. Trends in Cognitive Sciences, 15(2), 85–93. 10.1016/j.tics.2010.11.004

Gao, L., Cai, Y., Wang, H., Wang, G., Zhang, Q., & Yan, X. (2019). Probing prefrontal cortex hemodynamic alterations during facial emotion recognition for major depression disorder through functional near-infrared spectroscopy. Journal of Neural Engineering, 16(2), Article 026026. 10.1088/1741-2552/ab0093

García, A. S., Fernández-Sotos, P., Vicente-Querol, M. A., Lahera, G., Rodriguez-Jimenez, R., & Fernández-Caballero, A. (2020). Design of reliable virtual human facial expressions and validation by healthy people. Integrated Computer-Aided Engineering, 27(3), 287–299. 10.3233/ICA-200623

Garety, P. A., Edwards, C. J., Jafari, H., Emsley, R., Huckvale, M., Rus-Calafell, M., Fornells-Ambrojo, M., Gumley, A., Haddock, G., Bucci, S., McLeod, H. J., McDonnell, J., Clancy, M., Fitzsimmons, M., Ball, H., Montague, A., Xanidis, N., Hardy, A., Craig, T. K. J., & Ward, T. (2024). Digital AVATAR therapy for distressing voices in psychosis: The phase 2/3 AVATAR2 trial. Nature Medicine, 30(12), 3658–3668. 10.1038/s41591-024-03252-8

Gramfort, A., Luessi, M., Larson, E., Engemann, D. A., Strohmeier, D., Brodbeck, C., Goj, R., Jas, M., Brooks, T., Parkkonen, L., & Hämäläinen, M. (2013). MEG and EEG data analysis with MNE-Python. Frontiers in Neuroscience, 7, Article 267. 10.3389/fnins.2013.00267

Green, P., & MacLeod, C. J. (2016). SIMR: An R package for power analysis of generalized linear mixed models by simulation. Methods in Ecology and Evolution, 7(4), 493–498. 10.1111/2041-210X.12504

Hadley, L. V., Naylor, G., & Hamilton, A. F. D. C. (2022). A review of theories and methods in the science of face-to-face social interaction. Nature Reviews Psychology, 1(1), 42–54. 10.1038/s44159-021-00008-w

Hernandez, S. M., & Pollonini, L. (2020). NIRSplot: A tool for quality assessment of fNIRS scans. In Biophotonics Congress: Biomedical Optics 2020 (Translational, Microscopy, OCT, OTS, BRAIN) (Paper BM2C.5). Optica Publishing Group. 10.1364/BRAIN.2020.BM2C.5

Hortensius, R., Hekele, F., & Cross, E. S. (2018). The perception of emotion in artificial agents. IEEE Transactions on Cognitive and Developmental Systems, 10(4), 852–864. 10.1109/TCDS.2018.2826921

James, T. W., Potter, R. F., Lee, S., Kim, S., Stevenson, R. A., & Lang, A. (2015). How realistic should avatars be? An initial fMRI investigation of activation of the face perception network by real and animated faces. Journal of Media Psychology: Theories, Methods, and Applications, 27(3), 109–117. 10.1027/1864-1105/a000156

Joyal, C. C., Jacob, L., Cigna, M. H., Guay, J. P., & Renaud, P. (2014). Virtual faces expressing emotions: An initial concomitant and construct validity study. Frontiers in Human Neuroscience, 8, Article 787. 10.3389/fnhum.2014.00787

Ke, J., Song, H., Bai, Z., Rosenberg, M. D., & Leong, Y. C. (2025). Dynamic brain connectivity predicts emotional arousal during naturalistic movie-watching. PLOS Computational Biology, 21(4), Article e1012994. 10.1371/journal.pcbi.1012994

Kegel, L. C., Brugger, P., Frühholz, S., Grunwald, T., Hilfiker, P., Kohnen, O., Loertscher, M. L., Mersch, D., Rey, A., Sollfrank, T., Steiger, B. K., Sternagel, J., Weber, M., & Jokeit, H. (2020). Dynamic human and avatar facial expressions elicit differential brain responses. Social Cognitive and Affective Neuroscience, 15(3), 303–317. 10.1093/scan/nsaa039

Kragel, P. A., & LaBar, K. S. (2016). Decoding the nature of emotion in the brain. Trends in Cognitive Sciences, 20(6), 444–455. 10.1016/j.tics.2016.03.011

Krumhuber, E. G., Tamarit, L., Roesch, E. B., & Scherer, K. R. (2012). FACSGen 2.0 animation software: Generating three-dimensional FACS-valid facial expressions for emotion research. Emotion, 12(2), 351–363. 10.1037/a0026632

Kyrlitsias, C., & Michael-Grigoriou, D. (2022). Social interaction with agents and avatars in immersive virtual environments: A survey. Frontiers in Virtual Reality, 2, Article 786665. 10.3389/frvir.2021.786665

Lewis, J. P., Anjyo, K., Rhee, T., Zhang, M., Pighin, F., & Deng, Z. (2014). Practice and theory of blendshape facial models. In S. Lefebvre & M. Spagnuolo (Eds.), Eurographics 2014 – State of the art reports. The Eurographics Association. 10.2312/egst.20141042

Lindquist, K. A., & Barrett, L. F. (2012). A functional architecture of the human brain: Emerging insights from the science of emotion. Trends in Cognitive Sciences, 16(11), 533–540. 10.1016/j.tics.2012.09.005

Lindquist, K. A., Wager, T. D., Kober, H., Bliss-Moreau, E., & Barrett, L. F. (2012). The brain basis of emotion: A meta-analytic review. Behavioral and Brain Sciences, 35(3), 121–143. 10.1017/S0140525X11000446

Livingstone, S. R., & Russo, F. A. (2018). The Ryerson Audio-Visual Database of Emotional Speech and Song (RAVDESS): A dynamic, multimodal set of facial and vocal expressions in North American English. PLOS ONE, 13(5), Article e0196391. 10.1371/journal.pone.0196391

Lolansen, C., Dean Marshall, N., Fisher, C. T. L., & Burleigh, T. L. (2025). A decade of avatars: A systematic review of recreational gaming as a mechanism for gender identity exploration and affirmation. International Journal of Transgender Health. Advance online publication. 10.1080/26895269.2025.2595471

Luke, R., Larson, E. D., Shader, M. J., Innes-Brown, H., Van Yper, L., Lee, A. K. C., Sowman, P. F., & McAlpine, D. (2021). Analysis methods for measuring passive auditory fNIRS responses generated by a block-design paradigm. Neurophotonics, 8(2), Article 025008. 10.1117/1.NPh.8.2.025008

Miller, E. J., Foo, Y. Z., Mewton, P., & Dawel, A. (2023). How do people respond to computer-generated versus human faces? A systematic review and meta-analyses. Computers in Human Behavior Reports, 10, Article 100283. 10.1016/j.chbr.2023.100283

Oliver, M. M., & Amengual Alcover, E. (2020). UIBVFED: Virtual facial expression dataset. PLOS ONE, 15(4), Article e0231266. 10.1371/journal.pone.0231266

Oosterhof, N. N., & Todorov, A. (2008). The functional basis of face evaluation. Proceedings of the National Academy of Sciences, 105(32), 11087–11092. 10.1073/pnas.0805664105

Pessoa, L. (2017). A network model of the emotional brain. Trends in Cognitive Sciences, 21(5), 357–371. 10.1016/j.tics.2017.03.002

Pollonini, L., Bortfeld, H., & Oghalai, J. S. (2016). PHOEBE: A method for real time mapping of optodes-scalp coupling in functional near-infrared spectroscopy. Biomedical Optics Express, 7(12), 5104–5119. 10.1364/BOE.7.005104

R Core Team. (2026). R: A language and environment for statistical computing (Version 4.5.2) [Computer software]. R Foundation for Statistical Computing. https://www.r-project.org/

Reddy, P., Izzetoglu, M., Shewokis, P. A., Sangobowale, M., Diaz-Arrastia, R., & Izzetoglu, K. (2021). Evaluation of fNIRS signal components elicited by cognitive and hypercapnic stimuli. Scientific Reports, 11(1), Article 23457. 10.1038/s41598-021-02076-7

Saarimäki, H., Gotsopoulos, A., Jääskeläinen, I. P., Lampinen, J., Vuilleumier, P., Hari, R., Sams, M., & Nummenmaa, L. (2016). Discrete neural signatures of basic emotions. Cerebral Cortex, 26(6), 2563–2573. 10.1093/cercor/bhv086

Satpute, A. B., & Lindquist, K. A. (2019). The default mode network’s role in discrete emotion. Trends in Cognitive Sciences, 23(10), 851–864. 10.1016/j.tics.2019.07.003

Schindler, S., Zell, E., Botsch, M., & Kissler, J. (2017). Differential effects of face-realism and emotion on event-related brain potentials and their implications for the uncanny valley theory. Scientific Reports, 7(1), Article 45003. 10.1038/srep45003

Seabold, S., & Perktold, J. (2010). Statsmodels: Econometric and statistical modeling with Python. In S. van der Walt & J. Millman (Eds.), Proceedings of the 9th Python in Science Conference (pp. 92–96). 10.25080/Majora-92bf1922-011

Singh, A. K., & Dan, I. (2006). Exploring the false discovery rate in multichannel NIRS. NeuroImage, 33(2), 542–549. 10.1016/j.neuroimage.2006.06.047

Snoek, L., Jack, R. E., Schyns, P. G., Garrod, O. G. B., Mittenbühler, M., Chen, C., Oosterwijk, S., & Scholte, H. S. (2023). Testing, explaining, and exploring models of facial expressions of emotions. Science Advances, 9(6), Article eabq8421. 10.1126/sciadv.abq8421

Sollfrank, T., Kohnen, O., Hilfiker, P., Kegel, L. C., Jokeit, H., Brugger, P., Loertscher, M. L., Rey, A., Mersch, D., Sternagel, J., Weber, M., & Grunwald, T. (2021). The effects of dynamic and static emotional facial expressions of humans and their avatars on the EEG: An ERP and ERD/ERS study. Frontiers in Neuroscience, 15, Article 651044. 10.3389/fnins.2021.651044

Tupak, S. V., Dresler, T., Guhn, A., Ehlis, A. C., Fallgatter, A. J., Pauli, P., & Herrmann, M. J. (2014). Implicit emotion regulation in the presence of threat: Neural and autonomic correlates. NeuroImage, 85, 372–379. 10.1016/j.neuroimage.2013.09.066

Westgarth, M. M. P., Hogan, C. A., Neumann, D. L., & Shum, D. H. K. (2021). A systematic review of studies that used NIRS to measure neural activation during emotion processing in healthy individuals. Social Cognitive and Affective Neuroscience, 16(4), 345–369. 10.1093/scan/nsab017

Xu, G., Zhang, M., Wang, Y., Liu, Z., Huo, C., Li, Z., & Huo, M. (2017). Functional connectivity analysis of distracted drivers based on the wavelet phase coherence of functional near-infrared spectroscopy signals. PLOS ONE, 12(11), Article e0188329. 10.1371/journal.pone.0188329

