## Supplementary Materials for "Real versus virtual faces: Differentiating neural responses to emotional expressions"

Department of Psychology,

### Experiment 1

#### Methods

##### Stimulus

The complete set of 140 facial images used in Experiments 1 and 2 are presented in Fig S1-S4, with Matched-model images occurring in adjacent rows. Stimuli consisted of 20 unique face models - 10 drawn from the UIBVFED (5 male, 5 female), and 10 drawn from the RADIATE (5 male, 5 female). UIBVFED models were matched with RADIATE models on face shape, sex, skin tone, and hair colour. For each model, 7 facial expressions were then identified - Neutral, Joy, Sad, Angry, Fearful, Surprise, Disgust.

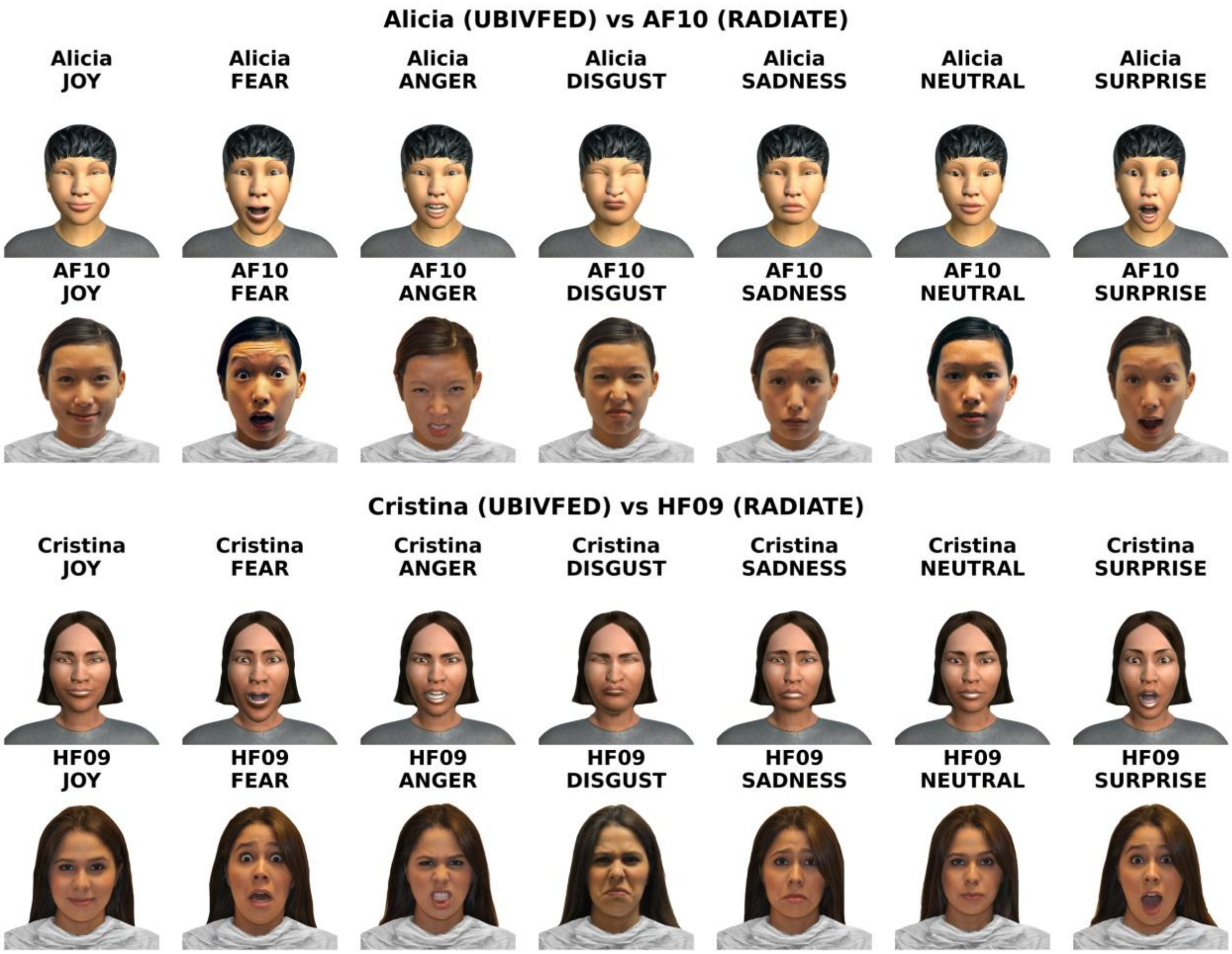


Figure S1: Matched-model stimulus from UBIVFED and RADIATE datasets, models 1-2.

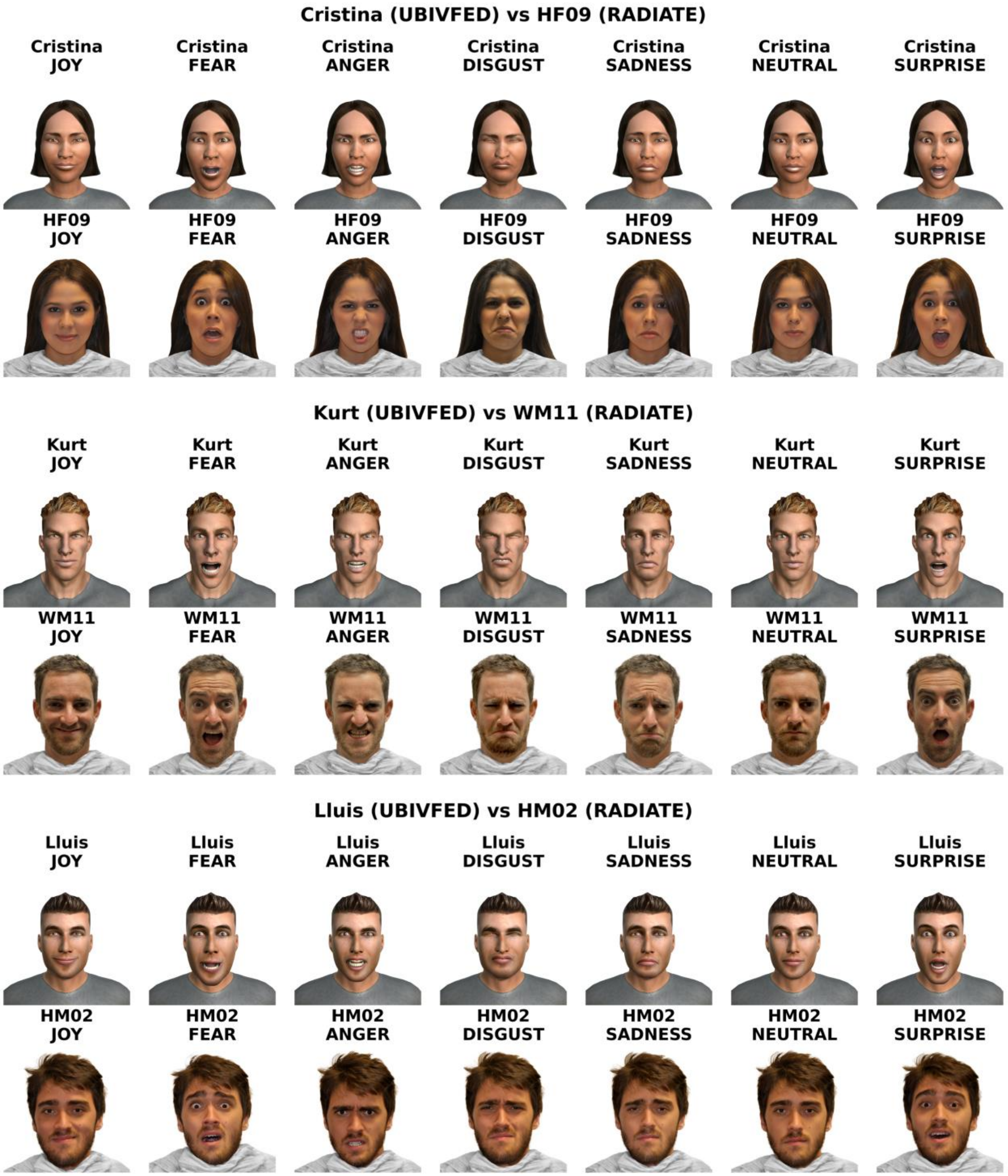


Figure S2: Matched-model stimulus from UBIVFED and RADIATE datasets, models 3-6.

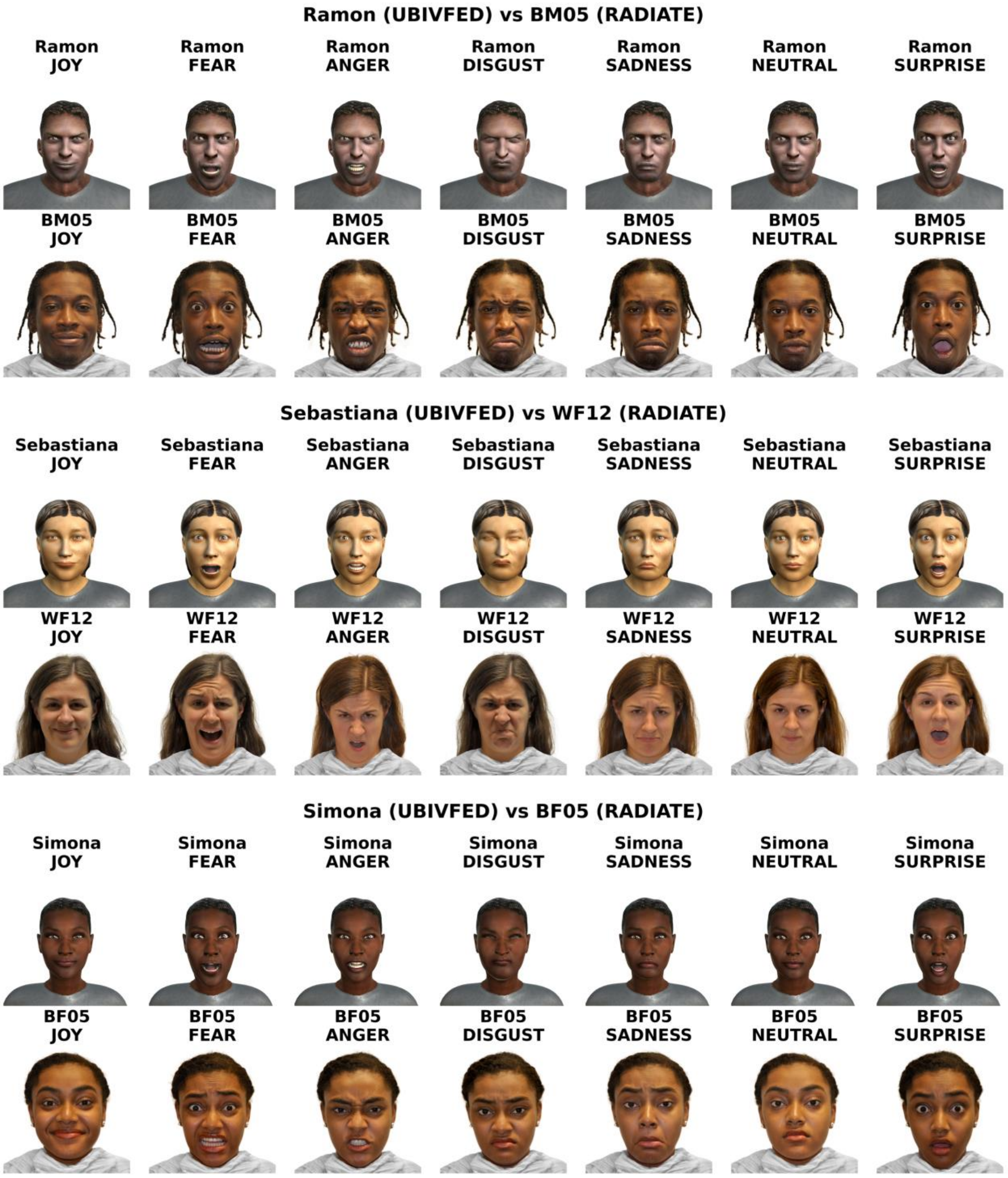


Figure S3: Matched-model stimulus from UBIVFED and RADIATE datasets, models 7-8.

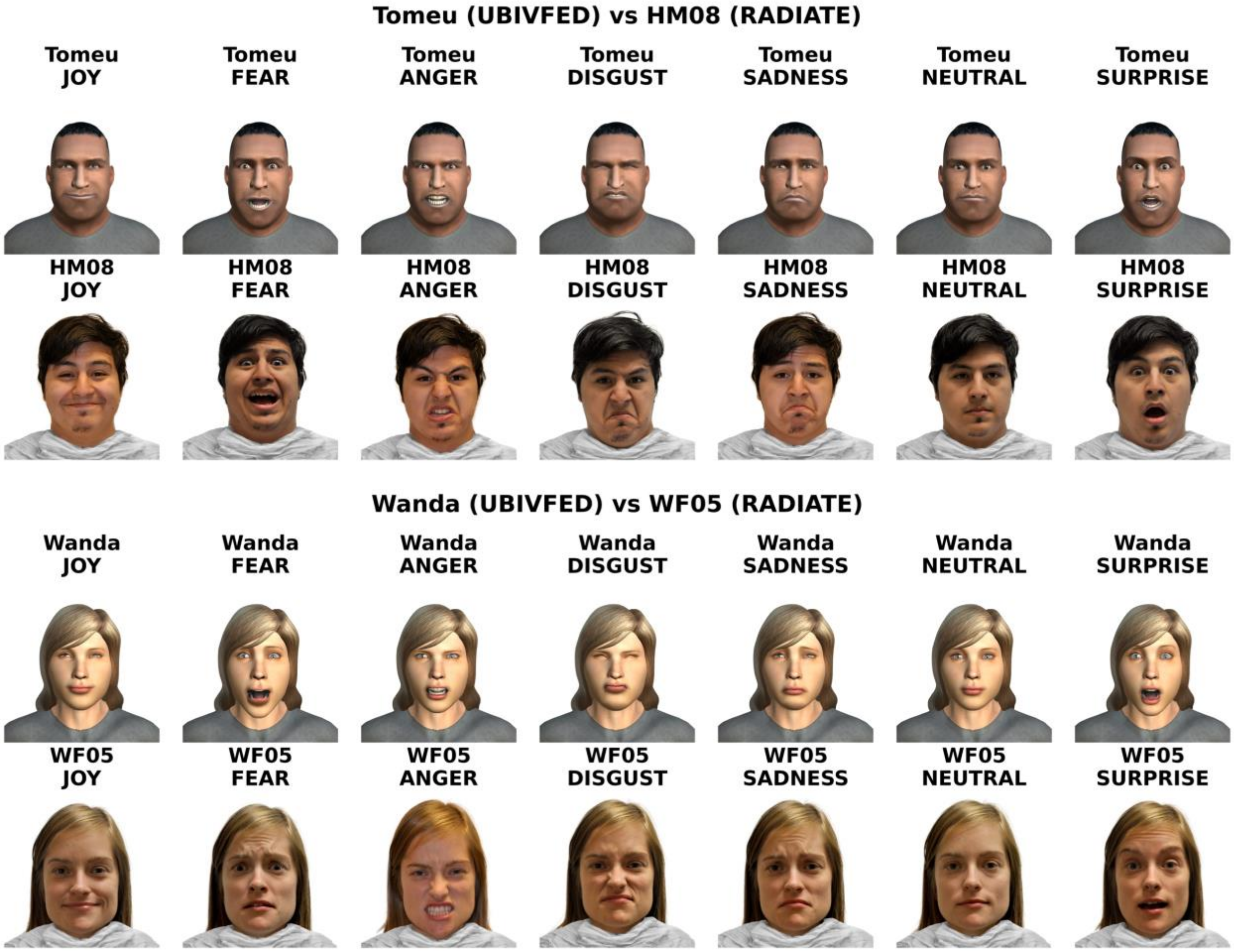


Figure S4: Matched-model stimulus from UBIVFED and RADIATE datasets, models 9-10.

#### Analyses

Data wrangling was performed using the "Tidyverse" package suite (Wickham et al., 2019). Similarity ratings were analysed with a mixed effects model using the "afex" package with afex::mixed() (Singmann et al., 2026). This method estimates the model with the "lme4" package and calculates test statistics with the "lmerTest" package (Bates et al., 2014; Kuznetsova et al., 2017). Test statistics, degrees of freedom, and p-value estimates were calculated using Kenward-Roger's approximation, as per current best practises (Luke, 2017), with restricted maximum likelihood (REML), fitted with the bobyqa optimizer (iterations = $1e^9$). Estimated marginal means were generated using the "emmeans" package, as were pairwise comparisons with Holm corrected p-values (Lenth et al., 2018). Effect size for the overall model was calculated as a conditional coefficient of determination, along with their 95% [CI], generated with 1000 parametric bootstrap iterations, using the package "partR2" (Stoffel et al., 2021). Figures were generated with additions from the "ggsignif" package (Ahlmann-Eltze & Patil, 2021).

#### Model building

We began by assessing the need for a mixed effects model. An intercept-only linear model fit with the gls function, and a random intercept model `(~1|participant)` fit with the lme function and maximum likelihood estimation (ML) using the "nlme" package (Pinheiro et al., 2007), were assessed with a likelihood ratio test. Once confirmed, we then fit a maximal fixed effects structure that was theoretically motivated (Zuur et al., 2009), which included all main effects and interactions for Congruence and Emotion. For the random effects structure, we then fitted the maximal random effects structure permitted by the experimental design to reduce instances of Type I errors (Barr et al., 2013). When the maximal model failed to converge, we reduced the random effects structure until convergence was achieved, and until the removal of additional structure resulted in a loss of power, as assessed with a LRT, as per Matuschek et al. (2017).

#### Results

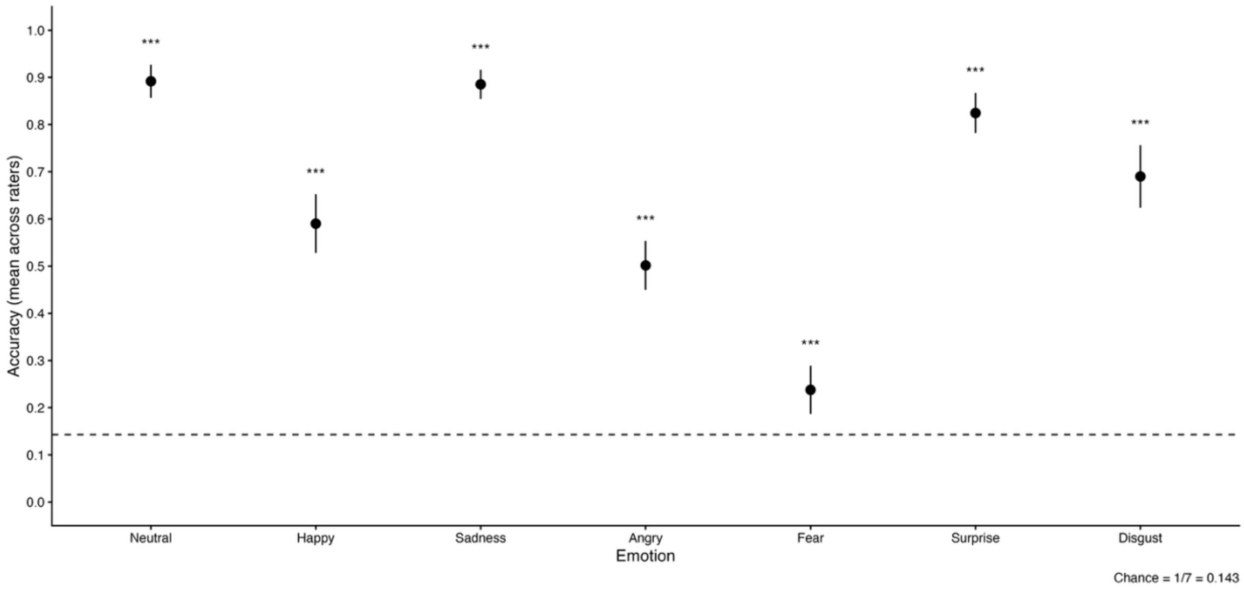


Figure S5: Mean emotion-identification accuracy (proportion correct) for each target emotion in the UBIVFED validation task. Points show mean per-rater accuracy with 95% confidence intervals; the dashed horizontal line indicates chance performance ($p_0 = {}^1/_7$). Asterisks denote Holm-adjusted one-sided tests against chance: * p <.05, ** p< .01, *** p<.001.

Experiment 2

### Results

Additional neuroimaging figures are presented below.

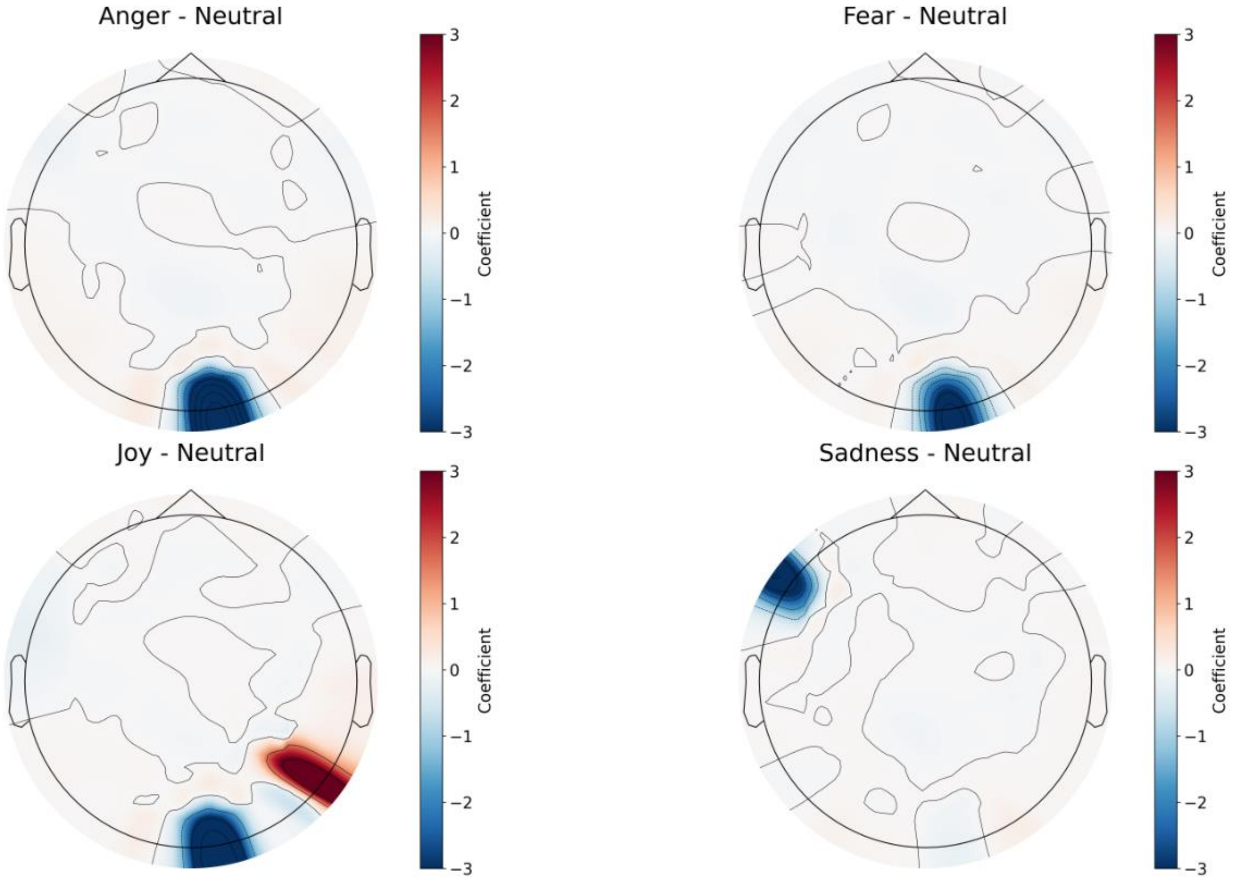


Figure S6: GLM results for the contrast between different emotions and neutral condition.

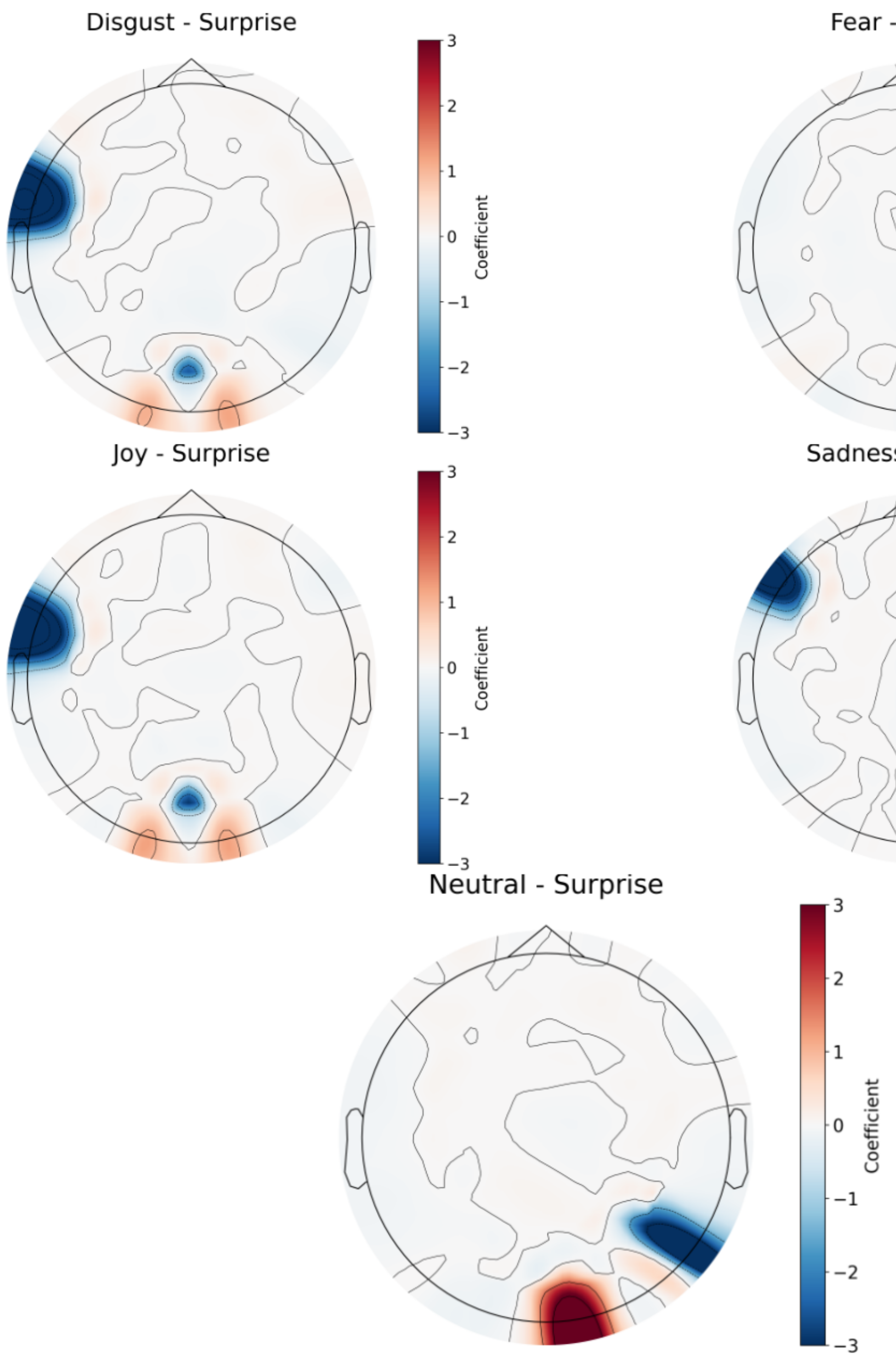


Figure S7: GLM results for the contrast between different emotions and surprise condition.

GLM Results for the Interaction of face type and emotion

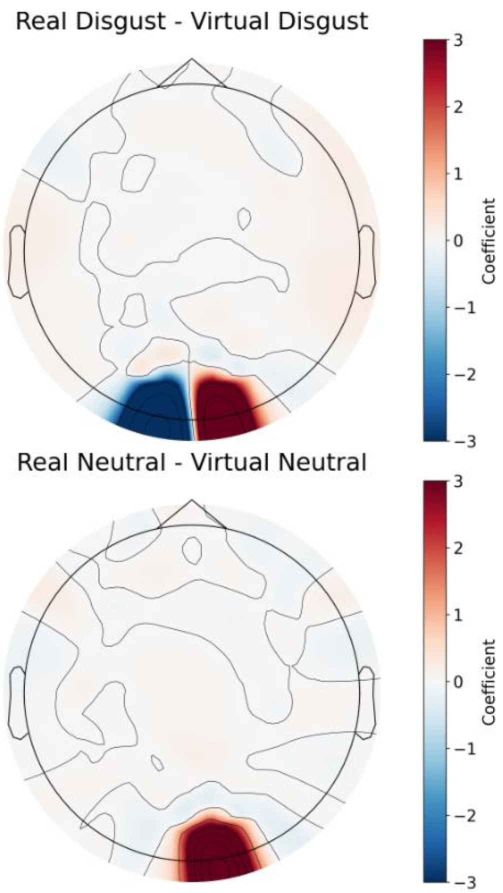


Figure S8: GLM results for the contrast between real and virtual conditions within each emotion.

Functional Connectivity contrasts for emotion

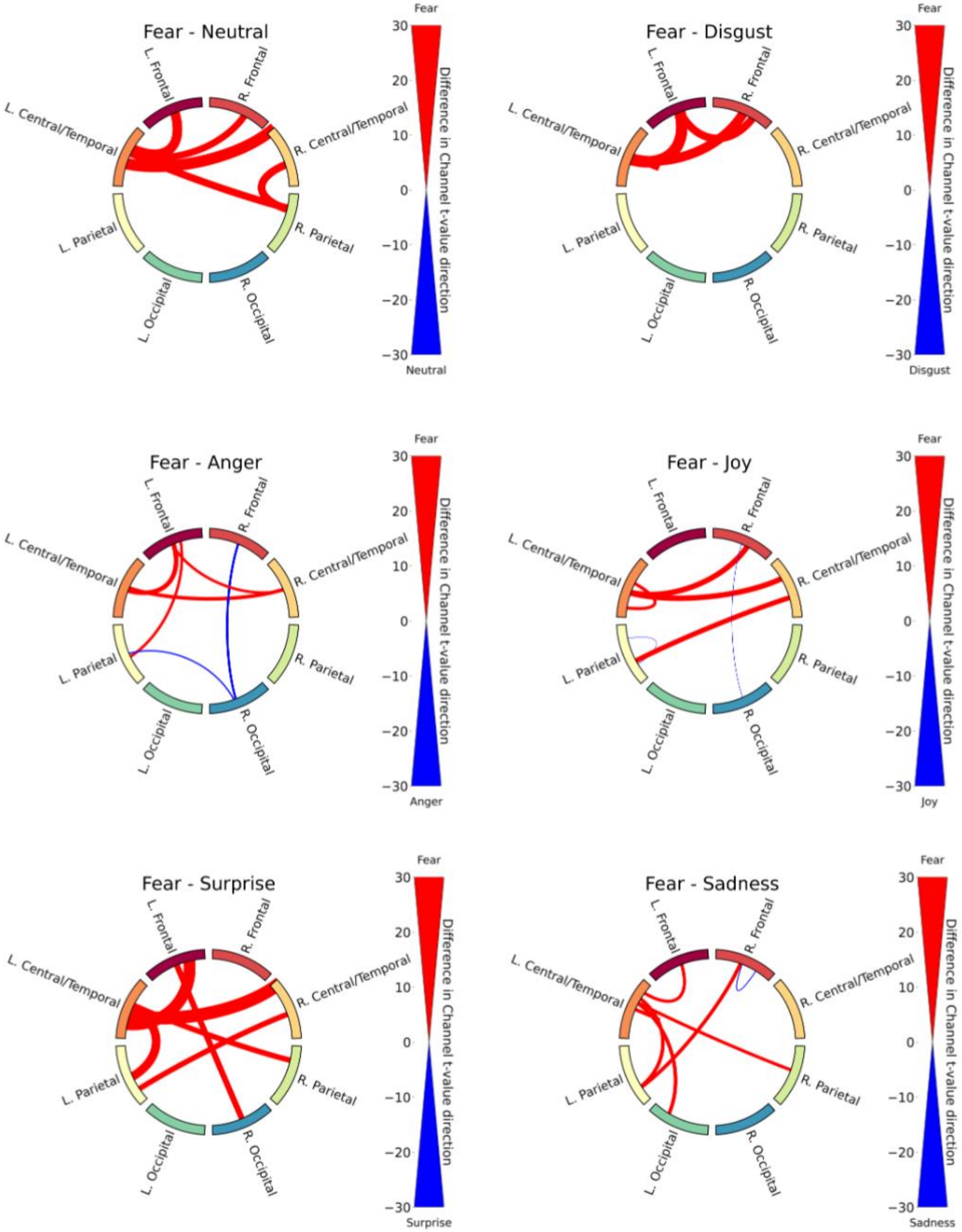


Figure S9: Functional connectivity results for emotion contrasts (1/4).

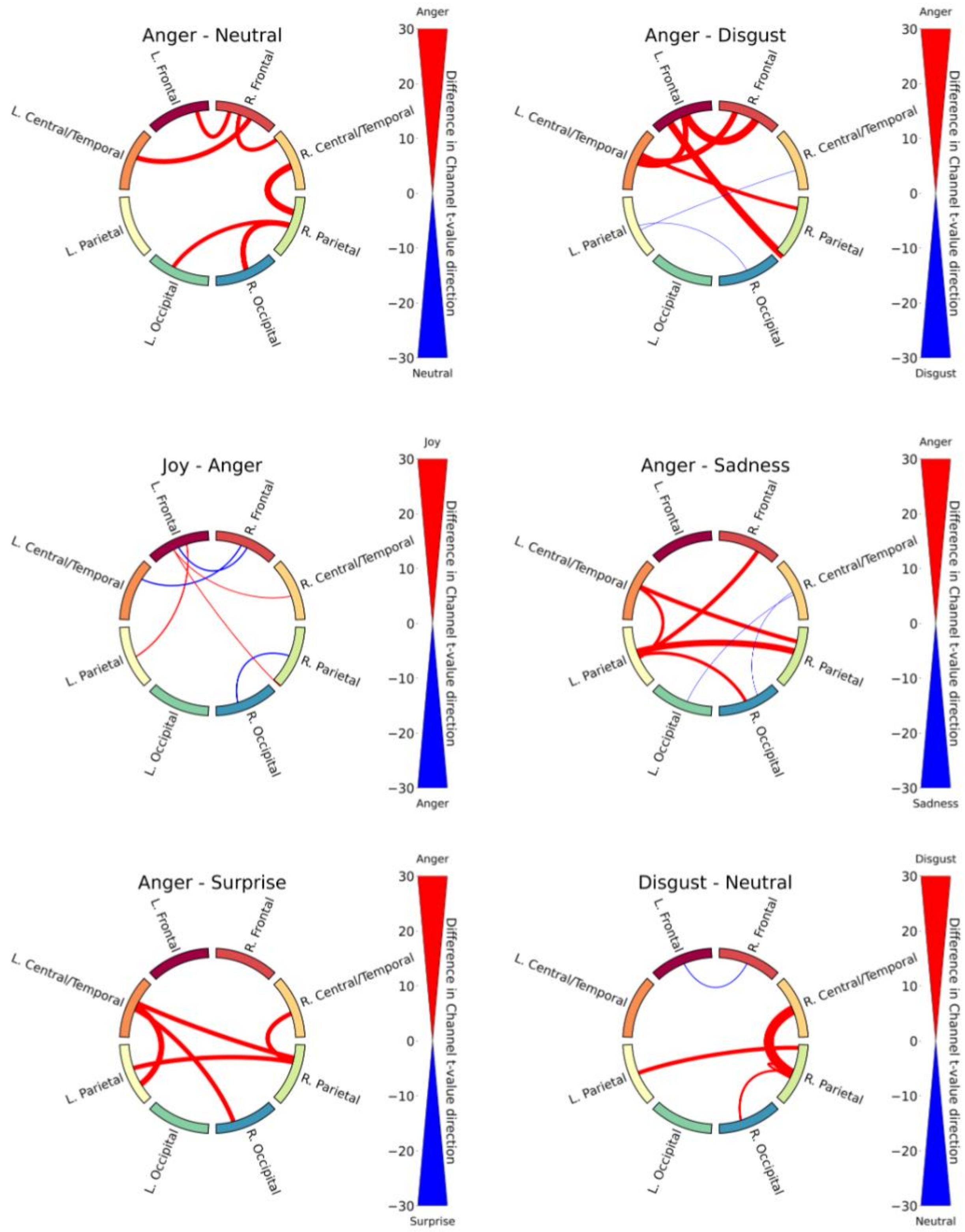


Figure S10: Functional connectivity results for emotion contrasts (2/4).

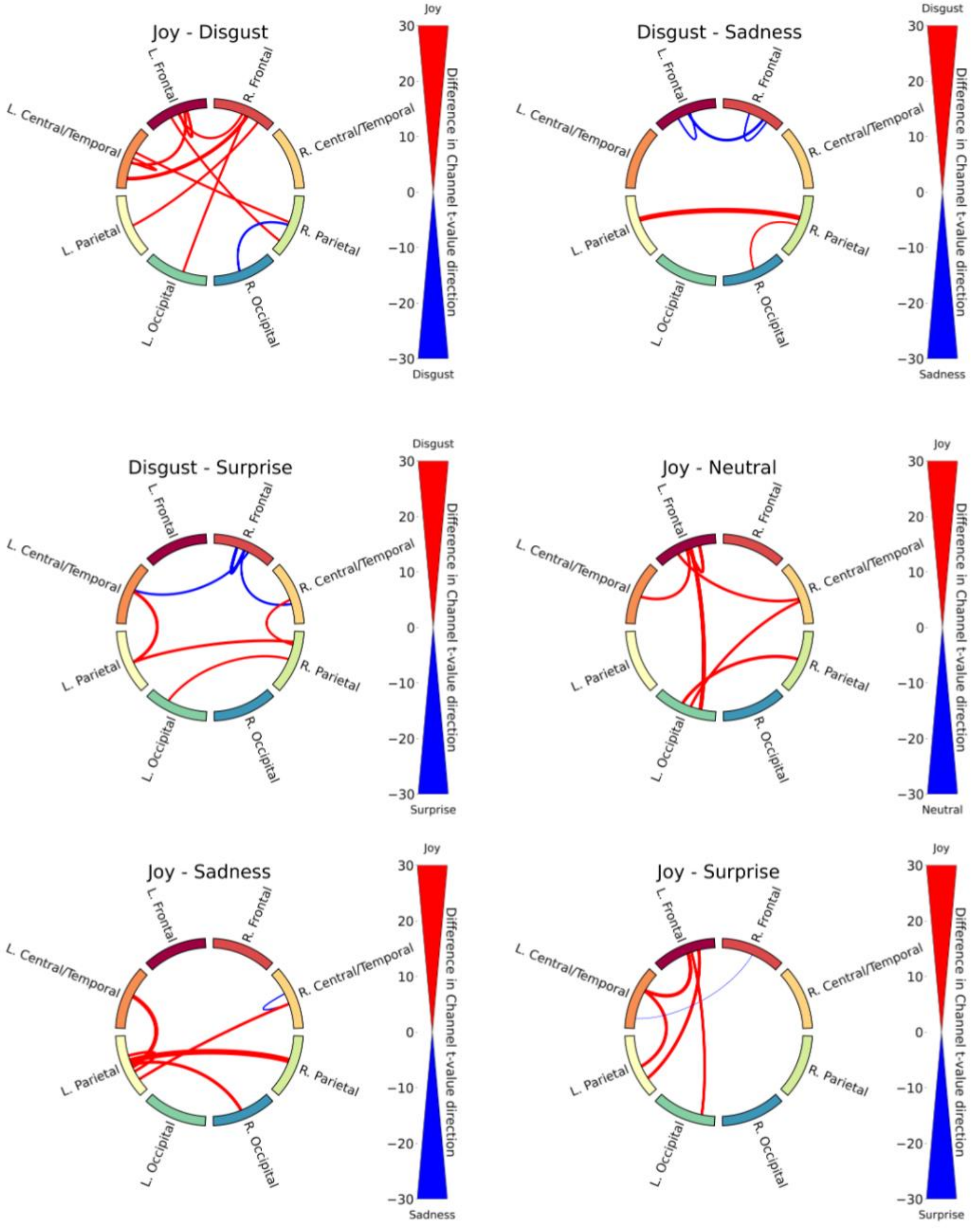


Figure S11: Functional connectivity results for emotion contrasts (3/4).

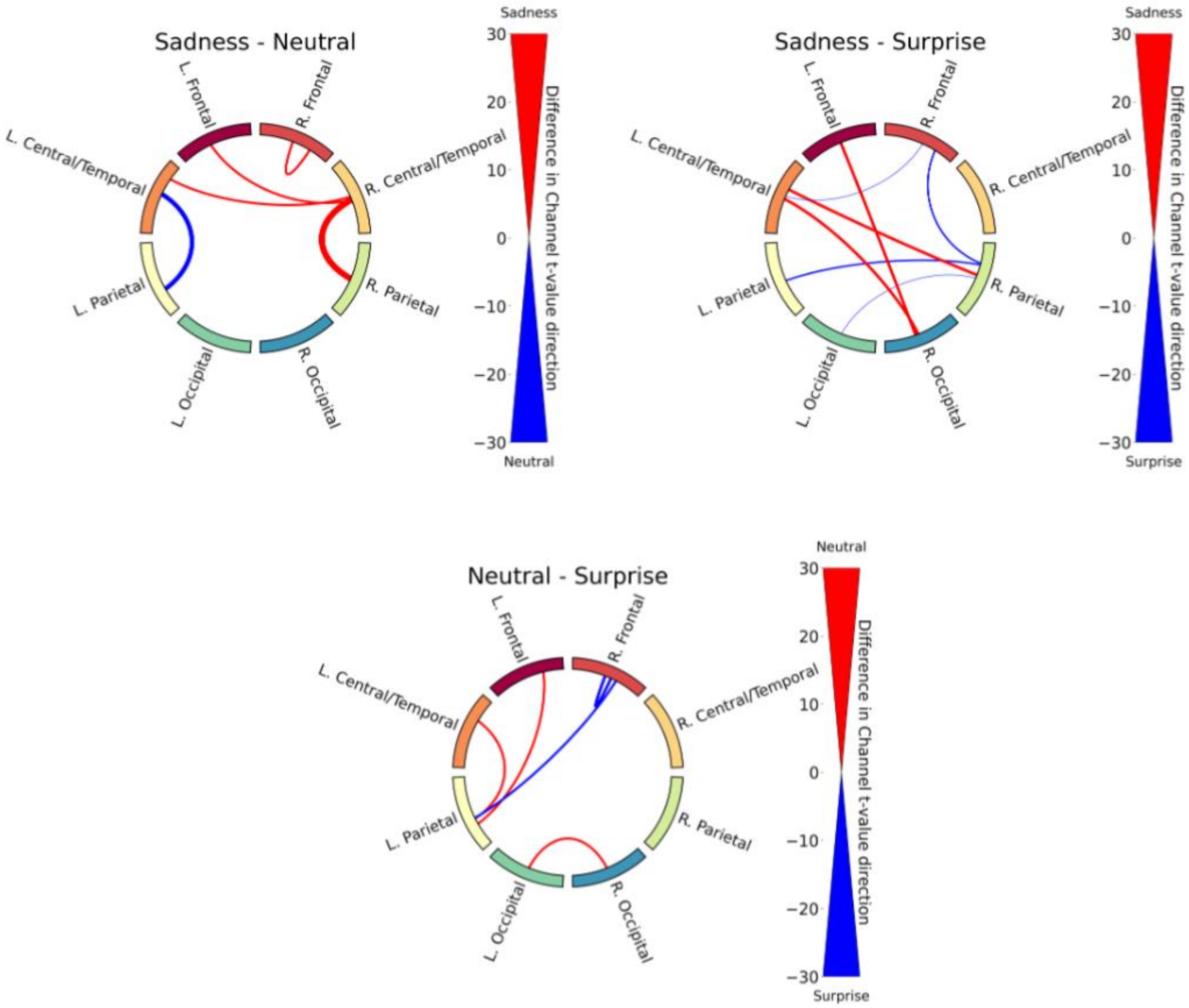


Figure S12: Functional connectivity results for emotion contrasts (4/4).

Functional Connectivity contrasts for the Interaction of face type and emotion

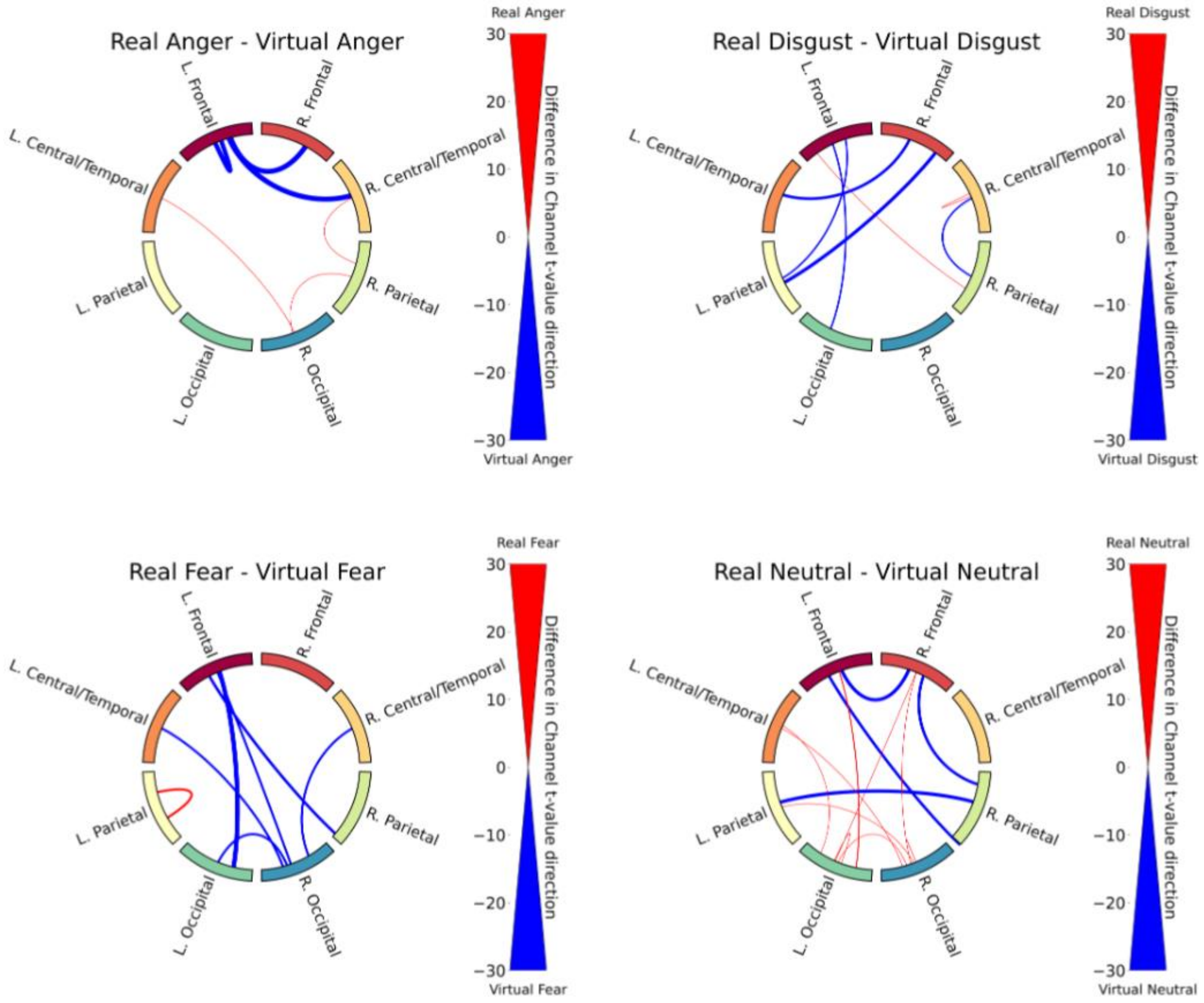


Figure S13: Functional connectivity results for the contrast between real and virtual conditions within each emotion. (1/2)

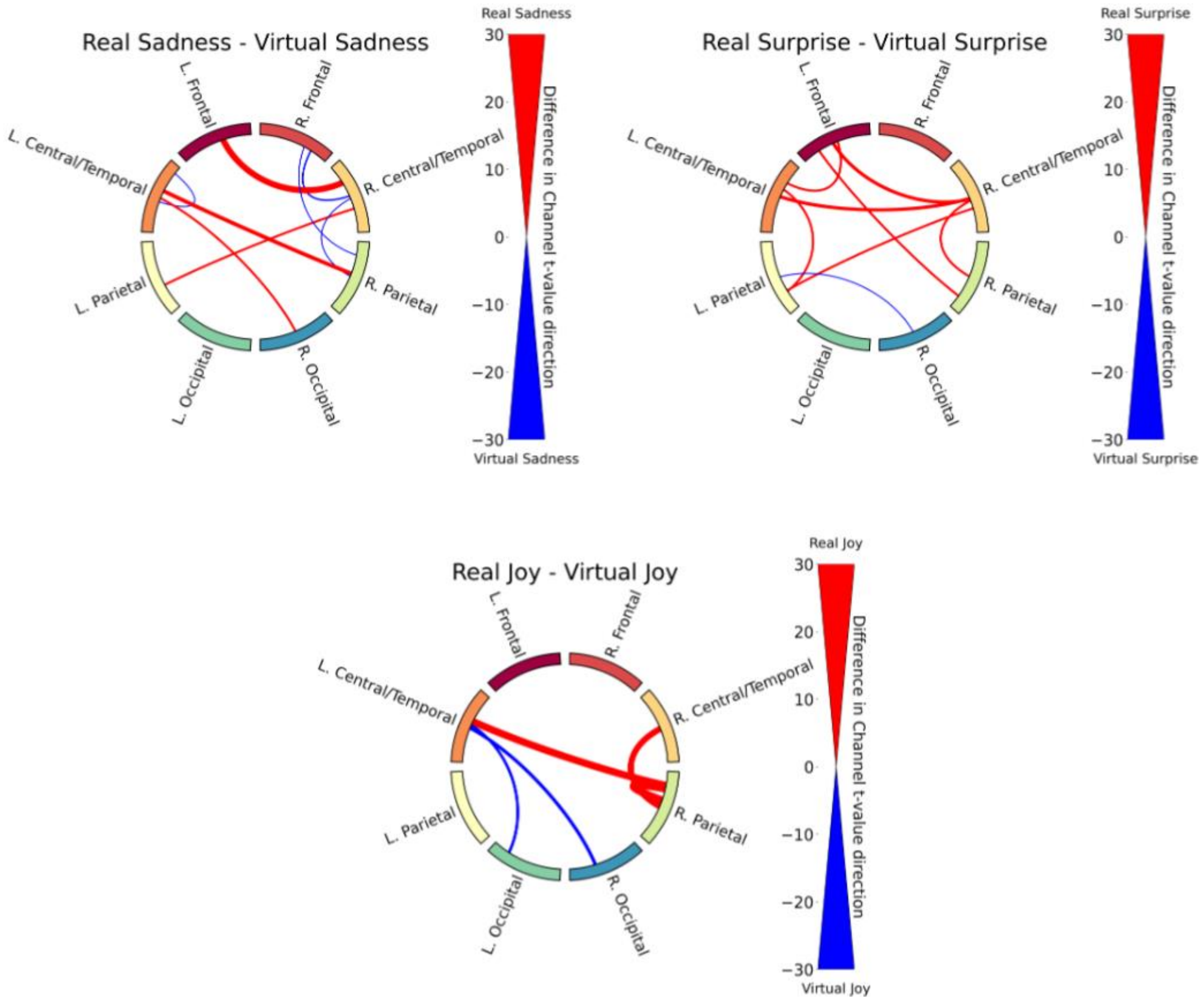


Figure S14: Functional connectivity results for the contrast between real and virtual conditions within each emotion. (2/2)

#### References

Ahlmann-Eltze, C., & Patil, I. (2021). *ggsignif: R package for displaying significance brackets for 'ggplot2'* [Preprint]. PsyArXiv. https://doi.org/10.31234/osf.io/7awm6

Barr, D. J., Levy, R., Scheepers, C., & Tily, H. J. (2013). Random effects structure for confirmatory hypothesis testing: Keep it maximal. *Journal of Memory and Language, **68***(3), 255–278. https://doi.org/10.1016/j.jml.2012.11.001

Bates, D., Mächler, M., Bolker, B., & Walker, S. (2014). *Fitting linear mixed-effects models using lme4* [Preprint]. arXiv. https://arxiv.org/abs/1406.5823

Kuznetsova, A., Brockhoff, P. B., & Christensen, R. H. B. (2017). lmerTest package: Tests in linear mixed effects models. *Journal of Statistical Software, **82***(13), 1–26. https://doi.org/10.18637/jss.v082.i13

Lenth, R., Singmann, H., Love, J., Buerkner, P., & Herve, M. (2018). *emmeans: Estimated marginal means, aka least-squares means* [Computer software]. https://CRAN.R-project.org/package=emmeans

Luke, S. G. (2017). Evaluating significance in linear mixed-effects models in R. *Behavior Research Methods, **49***(4), 1494–1502. https://doi.org/10.3758/s13428-016-0809-y

Matuschek, H., Kliegl, R., Vasishth, S., Baayen, H., & Bates, D. (2017). Balancing Type I error and power in linear mixed models. *Journal of Memory and Language, **94***, 305–315. https://doi.org/10.1016/j.jml.2017.01.001

Pinheiro, J., Bates, D., DebRoy, S., Sarkar, D., & R Core Team. (2007). *nlme: Linear and nonlinear mixed effects models* [Computer software]. https://CRAN.R-project.org/package=nlme

Singmann, H., Bolker, B., Westfall, J., Aust, F., & Ben-Shachar, M. S. (2026). *afex: Analysis of factorial experiments* (Version 1.5-1) [Computer software]. https://CRAN.R-project.org/package=afex

Stoffel, M. A., Nakagawa, S., & Schielzeth, H. (2021). partR2: Partitioning $R^2$ in generalized linear mixed models. *PeerJ, 9*, Article e11414. https://doi.org/10.7717/peerj.11414

Wickham, H., Averick, M., Bryan, J., Chang, W., McGowan, L. D., François, R., Grolemund, G., Hayes, A., Henry, L., Hester, J., Kuhn, M., Pedersen, T. L., Miller, E., Bache, S. M., Müller, K., Ooms, J., Robinson, D., Seidel, D. P., Spinu, V., . . . Yutani, H. (2019). Welcome to the Tidyverse. *Journal of Open Source Software, 4*(43), Article 1686. https://doi.org/10.21105/joss.01686

Zuur, A. F., Ieno, E. N., Walker, N. J., Saveliev, A. A., & Smith, G. M. (2009). *Mixed effects models and extensions in ecology with R*. Springer. https://doi.org/10.1007/978-0-387-87458-6
